# *De novo* production of prenylnaringenin compounds by a metabolically engineered *Escherichia coli*

**DOI:** 10.1101/2025.05.09.653005

**Authors:** Daniela Gomes, Joana L. Rodrigues, Nigel S. Scrutton, Ligia R. Rodrigues

## Abstract

Prenylnaringenin (PN) compounds, namely 8-prenylnaringenin (8-PN), 3’-prenylnaringenin (3’-PN), and 6-prenylnaringenin (6-PN), are reported to have several interesting bioactivities. This study aimed to validate a biosynthetic pathway for *de novo* production of PN in *Escherichia coli*. A previously optimized *E. coli* chassis capable of efficiently *de novo* producing naringenin was used to evaluate eleven prenyltransferases (PTs) for the production of PN compounds. As PT reaction requires dimethylallyl pyrophosphate (DMAPP) as extended substrate that has limited availability inside the cells, clustered regularly interspaced short palindromic repeats (CRISPR) and CRISPR-associated protein 12a (Cas12a) (CRISPR-Cas12a) was used to construct ten boosted DMAPP-E*. coli* strains. All the PTs, in combination with the naringenin biosynthetic pathway, were tested in these strains. Experiments in 96-well deep well plates identified twelve strains capable of producing PN. *E. coli* M-PAR-121 with the integration of the 1-deoxy-D-xylulose-5-phosphate synthase (DXS) gene from *E. coli* (*Ec*DXS) into the *lacZ* locus of the genome (*E. coli* M-PAR-121:*Ec*DXS) expressing the soluble aromatic PT from *Streptomyces roseochromogenes* (CloQ) and the naringenin biosynthetic pathway was selected as the best producer strain. After optimizing the production media in shake flasks, 160.57 µM of 3’-PN, 4.4 µM of 6-PN, and 2.66 µM of 8-PN were obtained. The production was then evaluated at the bioreactor scale and 397.57 µM of 3’-PN (135.33 mg/L) and 25.61 µM of 6-PN (8.72 mg/L) were obtained. To the best of our knowledge, this work represents the first report of *de novo* production of PN compounds using *E. coli* as a chassis.

## 1. Introduction

Prenylflavonoids are characterized by the presence of a lipophilic prenyl side-chain in the flavonoid skeleton, which confers higher lipophilicity and solubility. This characteristic leads to improved bioactivity due to enhanced interaction with target proteins (Mukai, 2018; Wen et al., 2021). Prenylnaringenin (PN) compounds, namely 8-prenylnaringenin (8-PN), 6-prenylnaringenin (6-PN), and 3’-prenylnaringenin (3’-PN), are reported to have several interesting biological activities, such as anticancer, antiviral, anti-inflammatory and estrogenic (Frattaruolo et al., 2024, 2019; Hitzman et al., 2020; Štulíková et al., 2018). PN compounds are naturally produced in some plant species in trace amounts, which makes their extraction and further incorporation into pharmaceutical and nutraceutical compounds difficult (Štulíková et al., 2018). Using microorganisms as microbial cell factories can provide an interesting alternative for their production, as it is a more efficient, environmentally friendly, and is potentially a more cost-effective method. The microbial production of PN compounds from a simple carbon source such as glucose depends on the efficient expression of several enzymes **(Fig. 1)**.

**Fig. 1.**
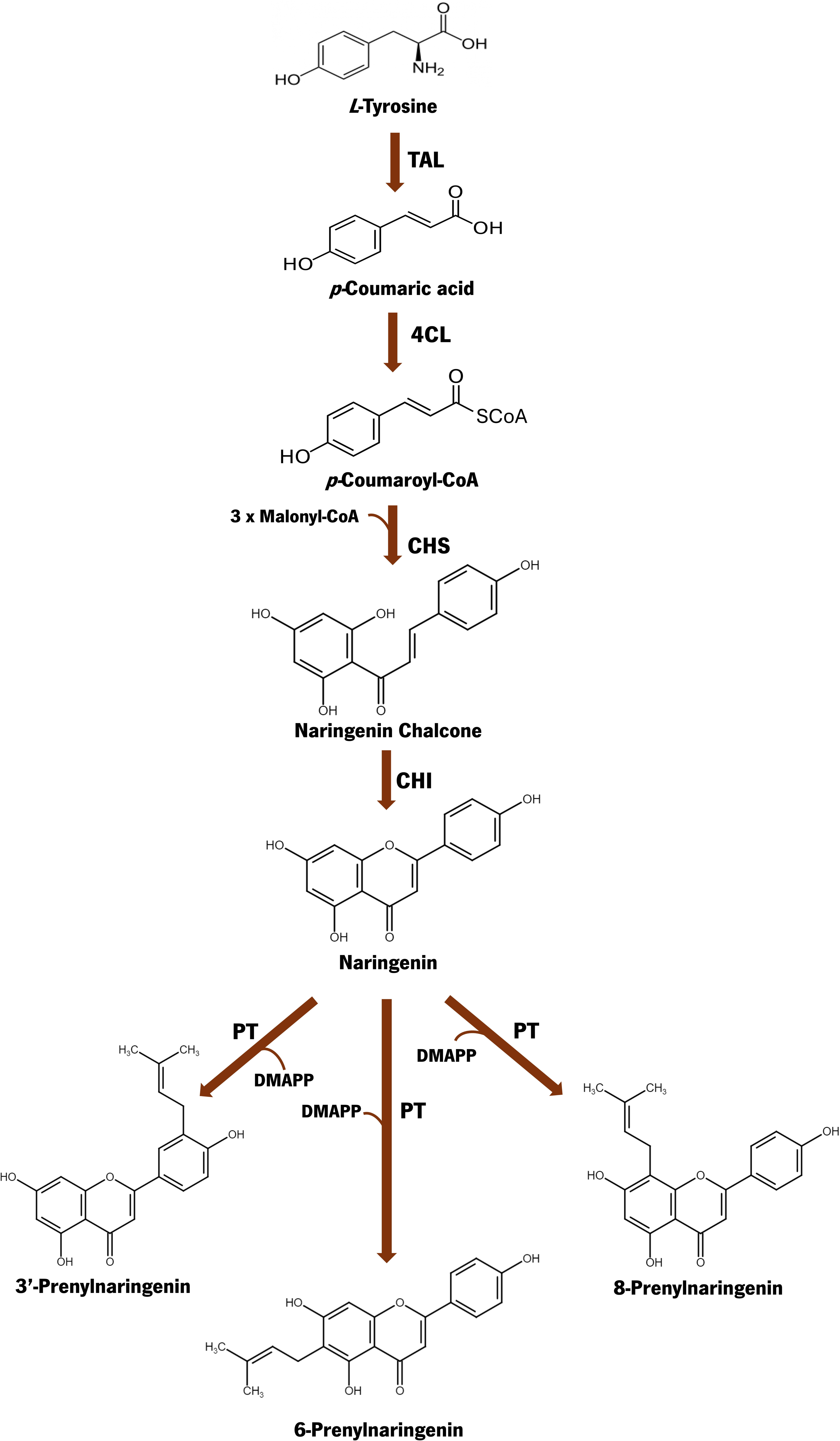

In a first step, the naringenin biosynthetic pathway should be assembled to produce the acceptor molecule in relevant amounts from glucose (Gomes et al., 2024). Afterwards, the prenylation step is catalyzed by a prenyltransferase (PT), that uses dimethyllallyl diphosphate (DMAPP) as extended substrate and transfers the prenylated chain to the acceptor molecule (naringenin). DMAPP is naturally produced in microorganisms through the mevalonate (MVA) or methylerythritol phosphate (MEP) pathways (Chatzivasileiou et al., 2019). The PT step can be considered the critical step for the production of prenylflavonoids since it depends on the limited DMAPP availability inside the cells and on the efficient expression of the PT enzymes (Peng et al., 2024).

Several aromatic PTs derived from plants or from microorganisms have been reported to perform prenylation in flavonoids (An et al., 2023; de Bruijn et al., 2020; Mori, 2020; Peng et al., 2024). Although PTs from plants are the only ones involved in the *in vivo* production of prenylflavonoids, these enzymes are membrane-bound enzymes, containing signal peptides that hinder their efficient expression in microorganisms. In contrast, microbial PTs are soluble enzymes making them easier to express and have been found to perform the prenylation of several flavonoid skeletons, including naringenin (de Bruijn et al., 2020; Peng et al., 2024).

In recent years, several efforts have been made to achieve the microbial production of prenylflavonoids. *Saccharomyces cerevisiae* was primarily exploited as chassis. *De novo* production of PN compounds in *S. cerevisiae* was first reported by Levisson et al. (2019). However, only 0.12 mg/L of 8-PN were produced. Later, Guo et al. (2022) reported the production of 49.35 mg/L and 101.40 mg/L of 8-PN from glucose in shake flask and 5 L bioreactor experiments, respectively (Guo et al., 2022). As an alternative to *S. cerevisiae*, *Escherichia coli* can also be explored as a chassis for the microbial production of PN compounds. The bioconversion of naringenin to PN compounds has been tested in *E. coli* using microbial PTs (Qiu et al. 2021; Liu et al. 2023; Zhang et al. 2024). More recently, *E. coli* was also engineered for the bioconversion of other flavonoids (silybin, daidzein, and baicalein) into their prenylated forms using fungal PTs (Fan et al. 2025). However, as far as we know, this microorganism has never been engineered to *de novo* produce prenylnaringenin or other prenylflavonoids.

In this work, we aimed to construct and validate a biosynthetic pathway to *de novo* produce PTs in *E. coli* for the first time. Using a previously optimized naringenin-producing *E. coli* strain, several PTs from plants and microbial sources were tested. Additionally, clustered regularly interspaced short palindromic repeats (CRISPR) and CRISPR-associated protein 12a (Cas12a) (CRISPR-Cas12a) was used to construct boosted DMAPP-*E. coli* strains to overcome this compound limitation. After screening all combinations of PTs/strains, the best producing strain was selected for production media optimization and bioreactor experiments. *E. coli* M-PAR-121 strain with the integration of the 1-deoxy-D-xylulose-5-phosphate synthase (DXS) gene from *E. coli* (*Ec*DXS) into the *lacZ* locus of the genome (*E. coli* M-PAR-121:*Ec*DXS) expressing the soluble aromatic PT from *Streptomyces roseochromogenes* (CloQ) and the naringenin biosynthetic pathway was able to produce 397.57 µM of 3’-PN (135.33 mg/L) and 25.61 µM of 6-PN (8.72 mg/L) at a bioreactor scale when two additional glucose pulses were provided, corresponding to the first report of *de novo* production in *E. coli* and the highest *de novo* production of PN reported in any host.

## 2. Materials and methods

### 2.1. Strains, plasmids, chemicals and media composition

*E. coli* NZY5α (NZYTech - MB00401) and *E. coli* NEB5α (New England Biolabs - C2987H) were used for cloning and for the propagation of plasmids. *E. coli* M-PAR-121 was used as the platform strain for the construction of DMAPP-modified strains (Koma et al., 2020). The heterologous biosynthetic pathways to produce PN were expressed in *E. coli* M-PAR-121 (wild-type strain) and in the *E. coli* M-PAR-121 DMAPP-modified strains. The features of all strains constructed and used in this work are presented in **Tab. S1**. The plasmids used in this study are presented in **Tab. S2**. Plasmids pWY16 and pWY24 were kindly provided by Dr. Shu-Ming Li (Yin et al. 2009; Yin et al. 2010). The prenyltransferases (PTs) carrying plasmids were already available in Scrutton laboratory. Plasmids pSIM*cpf1*, pTF-*lacZ*-*rfp*, and pBbS8c-*ddcpf1*-Δ, that were used in the clustered regularly interspaced short palindromic repeats (CRISPR)-Cas12 strategies, were designed and kindly provided by Jervis et al. (2021).

Isopropyl β-D-1-thiogalactopyranoside (IPTG), 5-bromo-4-chloro-3-indolyl-β-D-galactopyranoside (X-Gal) and lysogeny broth (LB) Miller medium were purchased from NZYTech. LB agar, used for colonies selection, was composed of 20 g/L LB Lennox (LabKem) and 20 g/L agar (LabKem). M9 minimal medium was composed by 3 g/L KH_2_PO_4_ (Riel-deHaën), 6 g/L Na_2_HPO_4_ (Chem-Lab), 0.5 g/L NaCl (NZYTech), 1 g/L NH_4_Cl (Panreac), 110 mg/L MgSO_4_ (Labkem), 15 mg/L CaCl_2_ (Panreac), 340 mg/L thiamine (Thermo Fisher Scientific), 5 g/L CaCO_3_ (Panreac), and vitamins (12.2 mg/L nicotinic acid (Acros organics), 10.8 mg/L pantothenic acid (Sigma Aldrich), 2.8 mg/L pyridoxine (Fisher BioReagents), 0.84 mg/L riboflavin (Panreac), 0.12 mg/L biotin (Merck), and 0.084 mg/L folic acid (Panreac)). Terrific broth (TB) was composed by 12 g/L tryptone (Fisher Scientific), 24 g/L yeast extract (LabKem), 9.4 g/L KH_2_PO_4_, and 2.2 g/L K_2_HPO_4_ (Panreac). M9 modified medium was composed by 5 g/L yeast extract (LabKem), 3 g/L KH_2_PO_4_, 6 g/L Na_2_HPO_4_ (Chem-Lab), 0.5 g/L NaCl, 1 g/L NH_4_Cl, 110 mg/L MgSO_4_ , 15 mg/L CaCl_2_ , 0.27 g/L FeCl_3_⋅6H_2_O (Panreac) and trace elements (0.2 g/L ZnCl_2_⋅4H_2_O (LabKem), 0.2 g/L CoCl_2_⋅6H_2_O (Sigma Aldrich), 0.2 g/L Na_2_MoO_4_⋅2H_2_O (Acros), 13 mg/L CuCl_2_⋅6H_2_O (Sigma Aldrich), 5 mg/L H_3_BO_3_ (Fisher Scientific), and 0.1 mL/L HCl (Fisher Scientific). M9 minimal medium, M9 modified medium, and TB medium were supplemented with glucose (Acros) at a final concentration of 30 g/L. The following antibiotics were used for strain selection: 50 µg/mL kanamycin (NZYTech), 25 µg/mL chloramphenicol (NZYTech), 100 µg/mL or 50 µg/mL spectinomycin (Alfa Aesar), 150 μg/mL hygromycin (Fischer Scientific).

### 2.2. Construction of the pathway plasmids

To construct a single plasmid carrying the complete naringenin biosynthetic pathway (composed by tyrosine-ammonia lyase (TAL) from *Flavobacterium johnsoniae* (*Fj*TAL), 4-coumarate:CoA ligase (4CL) from *Arabidopsis thaliana* (*At*4CL), chalcone synthase (CHS) from *Cucurbita maxima* (*Cm*CHS), and chalcone isomerase (CHI) from *Medicago sativa* (*Ms*CHI)), the cassette containing the *At*4CL and *Ms*CHI and respective promoters and terminators was amplified by polymerase chain reaction (PCR) using as template the previously constructed vector pACYCDuet_*At*4CL_*Ms*CHI (Gomes et al. 2024). The primers used for this PCR hold 15 bp homology overhangs for the pRSFDuet_*Fj*TAL_*Cm*CHS vector. pRSFDuet_*Fj*TAL_*Cm*CHS vector was also linearized by PCR. The construction of pRSFDuet_*Fj*TAL_*Cm*CHS_*At*4CL_*Ms*CHI was performed by In-Fusion cloning using the In-Fusion® Snap Assembly kit from Takara Bio Europe.

Eleven PTs were selected to be tested in this study (**Tab. S3**). PT from *Humulus lupulus* (*Hl*PT) and PT from *Sophora flavescens* (*Sf*N8DT-1) were synthesized with codon-optimization for *E. coli* by Twist Bioscience and then amplified by PCR. AnaPT and CdpC3PT from *Neosartorya fischeri* (without codon-optimization) were amplified by PCR from pWY16 and pWY24, respectively. These four PTs were cloned into the pCDFDuet-1 vector by restriction cloning. Codon-optimized versions of PT3 from *Cannabis sativa* (*Cs*PT3), coAnaPT, CloQ from *Streptomyces roseochromogenes,* PT from *E. coli* (*Ec*PT), NphB from *Streptomyces* sp., PT from *Streptomyces* sp. Act143 (*Sp*PT), and UbiA from *E. coli* were amplified by PCR from the respective carrying plasmids using primers with 15 bp homology overhangs for the pCDFDuet-1 vector. The pCDFDuet-1 vector was linearized by PCR and the cloning was performed by In-Fusion using the In-Fusion® Snap Assembly kit. The correct construction of these vectors was confirmed by colony PCR and sequencing. The primers (Metabion / Eurofins) used for vector linearization, genes amplification, colony PCR, and sequencing are displayed in **Tab. S4.**

### 2.3. Evaluation of PTs expression

*E. coli* M-PAR-121 carrying the PTs plasmids were grown at 37 °C in LB Miller, until reaching an optical density at 600 nm (OD_600nm_) of 0.6. The protein expression, the preparation of samples for sonication, the settings used for sonication, the preparation of protein samples and further quantification were previously described in Gomes et al. (2024). Soluble and insoluble fractions were subjected to sodium dodecyl sulfate (SDS) polyacrylamide gel electrophoresis (SDS-PAGE) gel (4% stacking gel and 10% running gel). Color Prestained Protein Standard, Broad Range (10-250 kDa) (NEB), Blue Prestained Protein Standard, Broad Range (11-250 kDa) (NEB), and NZYColour Protein Marker II (NZYTech) were used as reference protein ladders.

### 2.4. Construction of DMAPP-modified *E. coli* strains

DMAPP-modified *E. coli* strains were constructed sorting to the CRISPR-Cas12a system designed by Jervis et al. (2021).

#### 2.4.1. Genome integration of heterologous and native DXS and IDI genes

##### 2.4.1.1. Construction of target-specific integration vectors

To perform the genome integration of DXS and IDI genes, the pTF-*lacZ*-rfp vector previously constructed by Jervis et al. (2021) was used. This vector holds a CRISPR array and 500 bp upstream and downstream homology arms for the *lacZ* locus of the *E. coli* genome for efficient delivery and integration of cargo DNA. The red fluorescence protein coding gene (*rfp*), that was used as cargo DNA for integration in the study performed by Jervis et al. (2021), was removed by PCR using primers that anneal into the extremities of the *lacZ* arms. Linearized pTF-*lacZ* was used as backbone to receive the cassettes for genome integration.

Heterologous DXS gene from *Bacillus subtilis* (*Bs*DXS) and IDI genes from *S. cerevisiae* (*Sc*IDI) and *Bacillus licheniformis* (*Bl*IDI) were synthesized with codon-optimization for *E. coli* by Twist Bioscience. Native DXS and IDI from *E. coli* were amplified from *E. coli* M-PAR-121 genomic DNA extracted using Monarch Genomic DNA Purification Kit (NEB). The sequences of the native and heterologous codon optimized genes are listed in **Tab. S5.** Single and double integration of DXS and IDI genes with improved activities (*Bs*DXS, *Sc*IDI, and *Bl*IDI) were designed. Alternatively, single and double integration of the native DXS and IDI genes from *E. coli* were also designed. For integration of single genes, the genes were amplified by PCR with specific primers holding 15 bp homology arms for the linearized pTF-*lacZ* vector. In the case of double gene integration, the IDI gene was amplified with a forward primer with 15 bp homology for the corresponding DXS gene and a reverse primer with 15 bp homology for linearized pTF-*lacZ* vector and the DXS gene was amplified with a forward primer with 15 bp homology for linearized pTF-*lacZ* vector and a reverse primer that anneals in the end of the gene. The fragments were assembled by In-Fusion cloning using the In-Fusion® Snap Assembly kit. The following target-specific integration vectors were constructed: pTF-*lacZ-Bs*DXS, pTF-*lacZ-Sc*IDI, pTF-*lacZ*-*Bl*DI, pTF-*lacZ*-*Bs*DXS-*Sc*IDI, pTF-*lacZ*-*Bs*DXS-*Bl*IDI, pTF-*lacZ*-*Ec*DXS, pTF-*lacZ-Ec*IDI, and pTF-*lacZ*-*Ec*DXS-*Ec*IDI. The correct construction of these vectors was confirmed by colony PCR and sequencing. The primers (Metabion / Eurofins) used for vector linearization, genes amplification, colony PCR, and sequencing are displayed in **Tab. S6.**

##### 2.4.1.2. Clustered regularly interspaced short palindromic repeats (CRISPR) editing

CRISPR editing for the integration of the target-specific cassettes was performed as described by Jervis et al. (2021). *E. coli* M-PAR-121 chemically competent cells were transformed with pSIM*cpf1*. Then, electrocompetent cells of *E. coli* M-PAR-121 carrying pSIM*cpf1* were prepared and transformed with the constructed target-specific integration pTF-*lacZ* vectors. Recombinant colonies were selected at 30 °C in LB agar with 150 μg/mL of hygromycin and 50 μg/mL of spectinomycin. Since the cassette integration disrupted the *lacZ* gene, the blue-white screening test was performed by replating single colonies in LB agar containing the required antibiotics (hygromycin and spectinomycin), 0.1 mg/mL of X-GAL and 1 mM of IPTG. White colonies (positive integration recombinants) were selected and grown overnight at 30 °C in LB Miller with the required antibiotics. Afterwards, the genomic DNA was extracted using Monarch Genomic DNA Purification Kit (NEB). The correct integration into the *lacZ* locus was confirmed by PCR using the genomic DNA of each selected colony as template and by sequencing.

After confirmation, CRISPR plasmids were removed from the constructed strains as described by Jervis et al. (2021). Eight DMAPP-modified *E. coli* M-PAR-121 strains were constructed: *E. coli* M-PAR-121:*Bs*DXS, *E. coli* M-PAR-121:*Sc*IDI, *E. coli* M-PAR-121:*Bl*DI, *E. coli* M-PAR-121:*Bs*DXS-*Sc*IDI, and *E. coli* M-PAR-121:*Bs*DXS-*Bl*IDI, *E. coli* M-PAR-121:*Ec*DXS, *E. coli* M-PAR-121:*Ec*IDI, and *E. coli* M-PAR-121:*Ec*DXS-*Ec*IDI.

#### 2.4.2. Gene regulation using CRISPR interference (CRISPRi) system

##### 2.4.2.1. Construction of the CRISPRi vectors

Two strategies using the CRISPR interference (CRISPRi) system developed by Jervis et al. (2021) were designed to downregulate the geranyl diphosphate/farnesyl diphosphate synthase (*ispA*) gene. The protospacer adjacent motifs (PAM) TTTV, that is recognized by Cas12a for cleavage, were identified. Since targeting different PAM sequences of the gene can lead to differences in the downregulation levels, two different 23 bp array sequences were designed (Jervis et al., 2021). The first array was designed for the first PAM sequence (6 bp upstream of the *ispA* start codon) on the 5[ end of the gene coding sequence and the second array was designed for the PAM sequence 154 bp upstream of the *ispA* start codon. *ispA* gene with the PAM sequences and arrays highlighted are represented in **Tab. S7.** These arrays were introduced into the pBbS8c-*ddcpf1*-Δ by PCR using specific primers that were designed including the 23 bp array sequences and 15 bp homology arms for the vector to allow the vector circularization using the In-Fusion® Snap Assembly kit. The correct construction of pBbS8c-*ddcpf1*-*ispA*1 and pBbS8c-*ddcpf1*-*ispA*2 was confirmed by sequencing. The primers (Eurofins) used for vector linearization and insertion of the arrays and for sequencing are displayed in **Tab. S8.**

##### 2.4.2.2. Quantification of downregulation levels by reverse transcription-quantitative polymerase chain reaction (RT-qPCR)

*E. coli* M-PAR-121 was transformed with pBbS8c-*ddcpf1*-Δ (control), pBbS8c-*ddcpf1*-*ispA*1 and pBbS8c-*ddcpf1*-*ispA*2. Three colonies of each transformed strain were grown overnight in LB Miller medium supplemented with 25 µg/mL of chloramphenicol. These cultures were freshly cultivated (1:50) and grown at 37 °C until reaching an OD_600nm_ between 0.2-0.4. At this point, 10 mM *L-*arabinose and 0.1 mM IPTG were added for induction. The cells were cultivated at 37 °C and 200 rpm until reaching an OD_600nm_ between 1-1.4 and were then centrifuged to recover the pellet. The pellet was frozen in liquid nitrogen. Afterwards, the pellets were used for RNA extraction using the Direct-zol RNA Miniprep Kit (ZymoResearch) following the manufacturer’s instructions. RNA samples were run in 1% (w/v) agarose gel to check its integrity. The samples were quantified using Nanodrop One (Thermo). Afterwards, 1000 ng of RNA of each sample was used for cDNA synthesis. cDNA synthesis was performed using the iScript™ gDNA Clear cDNA Synthesis Kit (Bio-Rad) following the manufacturer’s instructions. Primers for reverse transcription-quantitative polymerase chain reaction (RT-qPCR) were designed using IDT Primer Quest Tool **(Tab. S9)**. The housekeeping genes putative 3-phenylpropionate transporter coding gene (*hcaT*) and gluconate transporter coding gene (*idnT*) were chosen as reference genes for normalization (Jervis et al., 2021; Zhou et al., 2011). RT-qPCR reaction was performed in technical triplicates for each sample using iTaq^TM^ Universal SYBR Green Supermix (Bio-Rad) in LightCycler® 480 System (Roche), following the manufacturer’s instructions.

### 2.5. Production experiments

Chemically competent cells of *E. coli* M-PAR-121 (wild-type (WT) strain) and DMAPP-modified *E. coli* M-PAR-121 strains were freshly prepared and transformed with the pathway plasmids using the heat-shock method. The transformants were selected in LB agar with the required antibiotics.

#### 2.5.1. 96-well deep well plates experiments

Overnight grown cultures of *E. coli* M-PAR-121 (WT strain) and DMAPP-modified *E. coli* M-PAR-121 strains transformed with the naringenin pathway plasmid (pRSFDuet_*Fj*TAL_*At*4CL_*Cm*CHS_*Ms*CHI) and the constructed PT plasmids were freshly cultivated (1:100) in 1 mL of TB medium supplemented with 4 g/L glucose and 100 µg/mL spectinomycin in 96-well deep well plates sealed with breathable sealing film. The 96-well deep well plate was incubated at 37 °C and 1000 rpm in a plate incubator shaker until reaching an OD_600nm_ between 1.5-2.0. At this point, IPTG was added at a final concentration of 0.1 mM and the temperature of incubation was changed to 30 °C. After 2 h, glucose was supplemented at a final concentration of 30 g/L to be used as substrate. The cultures were also maintained at 30 °C and 1000 rpm for 120 h. All the experiments were conducted in triplicate and samples were collected at the final time point for metabolites analysis.

#### 2.5.2. Shake flasks experiments

Overnight grown cultures of the producing strains carrying the naringenin pathway plasmid (pACYCDuet_*Fj*TAL_*At*4CL_*Cm*CHS_*Ms*CHI) and the PT plasmids selected in 96-well deep well plates experiments were used to inoculate 50 mL of LB Miller medium containing the required antibiotics in 250 mL shake flasks, at an initial OD_600nm_ of 0.1. The cultures were maintained at 37 °C and 200 rpm. IPTG (0.1 mM) was added when the culture attained an OD_600nm_ of 0.9 and the culture was then maintained at 26°C and 200 rpm for 5 h. Afterwards, a centrifugation was performed to pellet the cells (5000 rpm, 10 min) and the pellet was resuspended in M9 minimal medium supplemented with 30 g/L glucose and the relevant antibiotics. The experiment was maintained at 26 °C and 200 rpm for120 h (Gomes et al., 2024).

The best producers were also tested in single M9 modified and TB production media. Overnight grown cultures were used to inoculate 50 mL of M9 modified or TB media supplemented with 4 g/L glucose and the required antibiotics, at an initial OD_600nm_ of 0.1. The cultures were maintained at 37 °C and 200 rpm until attaining an OD_600nm_ of 0.9. Afterwards, 0.1 mM IPTG was supplemented, and the culture was maintained at 26°C and 200 rpm for 5 h. At this time, glucose at a final concentration of 30 g/L was supplemented to be used as substrate. The experiment was maintained at 26 °C and 200 rpm for 120 h. All the experiments were conducted in triplicate and samples were collected over time for metabolites analysis.

#### 2.5.3. Bioreactor experiments

Overnight grown culture of the best producing strain was used to inoculate 100 mL TB medium in 500 mL shake flasks at an initial OD_600nm_ of 0.1. This pre-culture was then incubated at 37 °C and 200 rpm for 12 h. Afterwards, the OD_600nm_ was evaluated and the volume of culture required to initiate the bioreactor experiments at an initial OD_600nm_ of 0.1 was centrifuged to pellet the cells at 5000 rpm for 10 min. The pellet was then resuspended in TB medium and used to inoculate the bioreactors. These experiments were performed in the 2L DASGIP® Parallel Bioreactor System (Eppendorf). All the bioreactor experiments were performed in duplicate and 2 mL samples were collected over time to measure OD_600nm_ and for metabolites analysis.

Batch experiments were performed in 400 mL of TB medium. Reactors were autoclaved at 121 °C for 20 min containing 350 mL of TB medium. The other 50 mL of TB medium were autoclaved separately in a shake flask and used to dissolve the cells for inoculation. An initial glucose concentration of 4 g/L and the required antibiotics were supplemented at the beginning of the experiment. The experiments were carried out at an initial temperature of 37 °C and initial agitation of 300 rpm, with a constant oxygen feeding of 0.5 vvm (12 L/h). The dissolved oxygen percentage (%DO) was maintained above 30% by adjusting stirring speed up to 1000 rpm. pH setting was 6.5 and the pH control was performed by automatic feeding a 2 M NaOH (LabSolve) solution. The cultures were maintained at 37 °C until the culture attained an OD_600nm_ of 0.9. At this OD_600nm_, IPTG at a final concentration of 0.1 mM was added to the reactor and the temperature setting was changed to 26 °C. After 5 h of induction, glucose was supplemented to the bioreactors at a final concentration of 30 g/L. The experiment was then maintained at 26 °C for 120 h.

Another batch experiment was performed in the same conditions. However, two additional pulses of glucose were supplemented during the experiment. Glucose concentrations were monitored over time using high-performance liquid chromatography (HPLC). Two pulses of glucose at a final concentration of 10 g/L were added to the bioreactors when the remaining glucose concentration on the medium was between 5-10 g/L (at 24 h and 63 h time points). The experiment was maintained at 26°C for 144 h.

### 2.6. Extraction of metabolites

#### 2.6.1. Methanol extraction

The metabolites produced in 96-well deep well plates experiments were extracted using 100% (v/v) methanol. An aliquot of the culture (100 µL) was transferred to a 96-well microplate and an equal volume of 100% (v/v) methanol was added. The 96-well microplate was vortexed for 2 min and then centrifuged for 10 min at 4000 rpm. The supernatant was recovered and added to a new 96-well microplate for ultra-performance liquid chromatography (UPLC) analysis.

#### 2.6.2. Ethyl acetate extraction

The metabolites produced in 50 mL shake flasks and bioreactor experiments were extracted using 100% (v/v) ethyl acetate. The extraction was performed from 1 mL whole broth culture in a 1:1 ratio. Samples extraction and preparation were performed as previously described in Gomes et al. (2024). Afterwards, the samples were analyzed by ultra-high performance liquid chromatography (UHPLC).

#### 2.6.3. Analytical methods

The metabolites produced in 96-well deep well plates experiments were evaluated by UPLC using the 1290 Infinity III Agilent LC system (Agilent, Santa Clara, United States) accoupled with a Kinetex® 5 µM XB-C18 100 Å LC column (50 x 2.1 mm). The diode array detector measured the absorbance at 290 nm. A binary mobile phase composed by 0.1% (v/v) formic acid in water (A) and acetonitrile (B) was used. The separation was achieved using a constant flow rate of 0.5 mL/min. The following elution gradient was used: 5% of B from 0-1 min, 5%-95% of B from 1-5 min, and 95-5% from 5-6 min. *p-*Coumaric acid, naringenin, 8-PN, 3’-PN, and 6-PN were quantified based on the peak areas at 1.0 min, 1.9 min, 2.8 min, 3.0 min, and 3.3 min, respectively. The metabolites produced in 50 mL shake flasks and bioreactor experiments were evaluated by UHPLC using the Shimadzu Nexera-X2 system (Shimadzu Corporation, Kyoto, Japan) accoupled with a Kinetex® 2.6 μm Polar C18 100 Å LC column (150 x 4.6 mm). SPD-M20A detector measured the absorbance at 290 nm. The binary mobile phase was also composed of 0.1% (v/v) formic acid in water (A) and acetonitrile (B), and the gradient was also maintained constant at a flow rate of 0.5 mL/min. The following gradient was used: 5%-95% of B from 1-10 min, 95%-5% of B from 10-12 min, and linearly 5% of B from 12-15 min. *p-*Coumaric acid, naringenin, 8-PN, 3’-PN, and 6-PN were quantified based on the peak areas at 7.5 min, 9.2 min, 10.6 min, 10.8 min, and 11.2 min, respectively.

Glucose consumption was evaluated in shake flasks and bioreactor experiments by HPLC sorting to a JASCO system connected with the RI-203 detector and a Aminex HPX-87 H column (Bio-Rad). The column was maintained at 60 °C. A constant flow rate of 0.5 mL/min of 5 mM H_2_SO_4_ was used, and glucose was detected and quantified based on the peak area at 10.9 min.

### 2.7. Statistical analysis

GraphPad Prism Software. Inc., version 8.0.1. was used to perform the statistical analysis of the results. All the results presented correspond to the mean value of three independent tests ± standard deviation (96-well deep well plate and shake flask experiments) or two independent tests ± standard deviation (bioreactor experiments). Statistical significance was evaluated using Ordinary one-way ANOVA tests. When *p-* value was <0.05, the differences were considered significant.

## 3. Results

### 3.1. Validation of PTs activity and their ability to produce PN compounds *de novo*

Functional PTs are required to produce PN compounds. In this study, eleven PTs from different sources were selected due to their reported activity to use flavonoids as acceptor substrates for the prenylation (Mori, 2020; Ozaki et al., 2009; Rea et al., 2019; Sasaki et al., 2011, 2009; Tello et al., 2008; Tsurumaru et al., 2012, 2010; Winkelblech et al., 2015; Zhou et al., 2015) Three PTs from plants were selected: PT from *H. lupulus* (*Hl*PT), N8DT-1 from *S. flavescens* (*Sf*N8DT-1), and PT3 from *C. sativa* (*Cs*PT3). Codon-optimized versions of the genes for *E. coli* were synthesized to avoid possible translation errors. However, plant PTs are membrane-bound enzymes being more difficult to be efficiently expressed in *E. coli* since this microbial chassis does not possess intracellular compartments and an endomembrane system similar to the ones present in plant cells. As an alternative, eight PTs from microbial sources were selected.

These microbial aromatic PTs are soluble enzymes being more easily expressed in *E. coli*. In two cases the original sequence without codon optimization was tested: CdpC3PT and AnaPT from *N. fischeri*. Moreover, codon-optimized versions of the following PTs were also tested: AnaPT from *N. fischeri* (coAnaPT), CloQ from *S. roseochromogenes,* PT from *E. coli (Ec*PT), NphB from *Streptomyces* sp., PT from *Streptomyces* sp. Act143 (*Sp*PT), and UbiA from *E. coli.* These PTs were cloned into the pCDFDuet-1 backbone that is widely used as expression vector for *E. coli* (Tolia et al. 2006). The constructed pCDFDuet_PT vectors were expressed in *E. coli* M-PAR-121, a tyrosine-overproducing strain constructed by Koma et al. (2020). *E. coli* M-PAR-121, *E. coli* K-12 MG1655 (DE3) and *E. coli* BL21 (DE3) were previously tested towards the production of naringenin. Higher productions of naringenin using glucose as substrate were obtained when *E. coli* M-PAR-121 was used as chassis (Gomes et al., 2024). Considering this, this strain was used as chassis to evaluate PN production. The expression of the eleven PT genes in *E. coli* M-PAR-121 was tested and evaluated by SDS-PAGE gel (**Fig. S1 and Fig. S2**). As expected, the SDS-PAGE gels for plant-derived PTs did not reveal the desired band for the proteins, indicating that these three PTs are not being efficiently expressed in *E. coli* or the amount of expressed protein is not enough to be visible on the gel. In contrast, the SDS-PAGE gels for aromatic PTs from microbial sources presented the desired protein bands indicating that these soluble enzymes are more easily expressed in *E. coli*.

Due to the relevance of producing PN compounds from a simple carbon source, the initial screening of the several PTs was performed directly from glucose by assembling the complete biosynthetic pathway in *E. coli* M-PAR-121. In order to reduce the possible metabolic burden of *E. coli* cells imposed by the expression of several plasmids, a single plasmid holding the previously optimized combination of genes of the naringenin pathway was constructed (pRSFDuet_*Fj*TAL_*Cm*CHS_*At*4CL_*Ms*CHI). The production of naringenin by *E. coli* M-PAR-121 expressing pRSFDuet_*Fj*TAL_*Cm*CHS_*At*4CL_*Ms*CHI was validated and 492.98 mg/L of naringenin were produced (**Fig. S3**). This strain was further transformed with the pCDFDuet_PT vectors and the constructed strains were further tested in 96-well deep well plate experiments using 30 g/L of glucose as substrate. However, the production of PN was not detected in any combination. This suggested that intracellular DMAPP availability was a limiting factor for PTs efficient activity posing a significant challenge for the successful production of prenylated compounds. Considering these results, the improvement of DMAPP availability inside of *E. coli* M-PAR-121 cells is essential to attain higher productions of PN compounds.

### 3.2. Construction of DMAPP-modified *Escherichia coli* strains

DMAPP is naturally synthesized in *E. coli* through the methylerythritol phosphate (MEP) pathway. However, this pathway contains rate-limiting steps impairing the production of DMAPP which limits its intracellular availability. Beyond this, DMAPP is also used in competing pathways to produce geranyl diphosphate (GPP), farnesyl diphosphate (FPP), and geranylgeranyl diphosphate (GGPP) and in the synthesis of terpenoids and sterols (Chatzivasileiou et al., 2019; Henry et al., 2018). Consequently, finding strategies to improve the pool of DMAPP is mandatory to construct an efficient strain able to produce prenylnaringenin.

Since DXS and IDI were identified as rate-limiting steps in the MEP pathway, we attempted to overexpress these genes to improve their flux. *Bs*DXS was selected to be integrated and overexpressed in the *E. coli* M-PAR-121 genome since it has previously shown positive effects in the production of terpenoids, lycopene and isoprene, that also require DMAPP as extender substrate, in engineered *E. coli* (Chen et al., 2013; Rinaldi et al., 2022; Zhao et al., 2011). Moreover, *Sc*IDI and IDI from *Bl*IDI were selected since these genes showed higher activity for the conversion of IPP into DMAPP (Gao et al., 2016). The single integration of these genes and also the combination of *Bs*DXS with each IDI gene into the *lacZ* locus of the genome was performed and confirmed by PCR and sequencing. The strains *E. coli* M-PAR-121:*Bs*DXS, *E. coli* M-PAR-121:*Sc*IDI, *E. coli* M-PAR-121:*Bl*DI, *E. coli* M-PAR-121:*Bs*DXS-*Sc*IDI, and *E. coli* M-PAR-121:*Bs*DXS-*Bl*IDI were successfully constructed using the CRISPR-Cas12a integration system (**Fig. S4**). Native DXS and IDI from *E. coli* were also overexpressed by integrating another gene copy into the *lacZ* locus of the genome. Single *Ec*DXS, single *Ec*IDI and a cassette holding both genes were integrated into the *E. coli* M-PAR-121 and the successful integration was confirmed by PCR and sequencing. The strains *E. coli* M-PAR-121:*Ec*DXS, *E. coli* M-PAR-121:*Ec*IDI, and *E. coli* M-PAR-121:*Ec*DXS-*Ec*IDI were successfully constructed using the CRISPR-Cas12a integration system (**Fig. S5**).

Two strategies for downregulation of the geranyl diphosphate/farnesyl diphosphate synthase (*ispA*) gene, responsible for DMAPP conversion into FPP, were also designed. Since *ispA* is considered an essential gene for *E. coli* due to its function in the isoprenoid pathway and production of essential lipids for cell wall maintenance, the complete gene knockout would probably result in cell death (Mendez-Perez et al., 2017; Shiomi and Niki, 2011). Two different arrays were designed to target two different PAM sequences of the gene using CRISPRi since it was already reported that targeting different PAM sequences can lead to differences in the downregulation levels. Moreover, it was reported that the gene repression level is higher the closer the PAM region is to the initiation translation codon (Jervis et al., 2021; Tao et al., 2018). The gene repression was evaluated by RT-qPCR by comparing the *ispA* expression levels in *E. coli* M-PAR-121 with pBbS8c-*ddcpf1*-*ispA*1 and *E. coli* M-PAR-121 with pBbS8c-*ddcpf1*-*ispA*2 with the expression levels of the control strain (*E. coli* M-PAR-121 with pBbS8c-ddcpf1-Δ) and two housekeeping genes (*hcaT* and *idnT).* The melting curves of the primers used for each target to ensure primer efficiency are represented in **Fig. S6**. After calculating the expression fold change, it was possible to conclude that *E. coli* M-PAR-121 with pBbS8c-*ddcpf1*-*ispA*1 (array targeting the first PAM sequence after the *ispA* start codon) leads to an *ispA* downregulation of 73%. As expected, *E. coli* M-PAR-121 with pBbS8c-*ddcpf1*-*ispA*2 with the array targeting a PAM sequence 154 bp upstream of the *ispA* start codon, leads to a lower level of *ispA* downregulation (46%).

### 3.3. Evaluation of *de novo* PN production in the constructed DMAPP-modified *E. coli* strains

#### 3.3.1. Screening in 96-well deep well plates experiments

The ten DMAPP-modified *E. coli* constructed strains were transformed with pRSFDuet_*Fj*TAL_*Cm*CHS_*At*4CL_*Ms*CHI and the pCDFduet-1 vectors holding the different PTs. One hundred and ten (110) combinations of strains/PTs were obtained to perform production experiments in 96-well deep well plates. Since the complete biosynthetic pathway to produce PN from a simple carbon source was constructed, only glucose (at a final concentration of 30 g/L) was supplemented to these experiments to be used as substrate. Out of the 110 combinations, only 12 were able to *de novo* produce PN from glucose (**Tab. 1**). 3’-PN was the most produced PN compound and 6-PN was also detected in two of the combinations.

**Tab. 1.** *De novo* production of prenylnaringenin (PN) compounds in 96-well deep well plates experiments. Production experiments were performed in triplicate for the 110 combinations of strains expressing the pRSFDuet_*Fj*TAL_*Cm*CHS_*At*4CL_*Ms*CHI (NAR plasmid) and the pCDFDuet vectors holding prenyltransferases (PTs). Only the 12 combinations able to *de novo* produce prenylnaringenin are herein represented. The production of *p-*coumaric acid (CA), naringenin (NAR), 3’-prenylnaringenin (3’-PN), and 6-prenylnaringenin (6-PN) was evaluated by ultra-high performance liquid chromatography (UHPLC).

| Strain | PT | Produced compound (μM) |  |  |  |
| --- | --- | --- | --- | --- | --- |
|  |  | CA | NAR | 3'-PN | 6-PN |
| *E. coli* M-PAR-121:*BsDXS* | CdpC3PT | 546.13 ± 14.20 | 44.70 ± 8.56 | 21.64 ± 2.27 | - |
|  | CoAnaPT | 539.29 ± 37.72 | 41.72 ± 6.45 | 19.55 ± 0.05 | - |
|  | CloQ | - | 40.62 ± 2.49 | - | 3.31 ± 0.04 |
|  | NphB | 150.62 ± 18.83 | 57.92 ± 5.26 | 24.66 ± 1.49 | - |
|  | *Sp*PT | 463.29 ± 10.34 | 57.55 ± 3.67 | 37.01 ± 2.34 | - |
|  | UbiA | 182.96 ± 0.99 | 13.64 ± 2.45 | 20.56 ± 1.08 | - |
| *E. coli* M-PAR-121:*EcDXS* | CoAnaPT | - | 15.49 ± 0.53 | 15.29 ± 0.73 | - |
|  | CloQ | 1427.79 ± 360.00 | 57.79 ± 18.38 | 88.34 ± 15.22 | 6.20 ± 1.20 |
|  | NphB | 50.29 ± 2.84 | 18.69 ± 0.30 | 27.46 ± 2.27 | - |
|  | *Sp*PT | 268.07 ± 9.44 | 33.50 ± 3.65 | 19.64 ± 0.48 | - |
| *E. coli* M-PAR-121:*BsDXS*:*ScIDI* | CloQ | - | 42.16 ± 2.67 | 20.64 ± 2.30 | - |
| *E. coli* M-PAR-121:*EcDXS*:*EcIDI* | *Ec*PT | 764.15 ± 80.78 | 84.47 ± 4.85 | 49.55 ± 1.57 | - |

In the other 98 combinations, no peak was detected for PN compounds. Moreover, PN production was not detected when the CRISPRi strains (*E. coli* M-PAR-121 with pBbS8c-*ddcpf1*-*ispA*1 and *E. coli* M-PAR-121 with pBbS8c-*ddcpf1*-*ispA*2) were used as microbial chassis. The production of PN was only detected in the *E. coli* M-PAR-121:*Bs*DXS, *E. coli* M-PAR-121:*Ec*DXS, *E. coli* M-PAR-121:*Bs*DXS:*Sc*IDI, and *E. coli* M-PAR-121:*Ec*DXS:*Ec*IDI. Among the tested PTs, PN compounds were not detected when plant aromatic PTs were expressed. The best producing strain in these experiments was *E. coli* M-PAR-121:*Ec*DXS expressing pRSFDuet_*Fj*TAL_*Cm*CHS_*At*4CL_*Ms*CHI and pCDFDuet_CloQ. This strain was able to produce 88.34 ± 15.22 µM of 3’-PN and 6.20 ± 1.20 µM of 6-PN. In addition to PN production, higher amounts of *p-*coumaric acid and naringenin were produced and accumulated in this strain compared with the other producing strains.

#### 3.3.2. Evaluation of the identified producing strains in shake flask experiments

The ability of the 12 identified strains to *de novo* produce PN was further evaluated in shake flask experiments. The production experiments were performed in a two-step approach using the combination of LB Miller and M9 minimal media supplemented with 30 g/L of glucose. The first step using LB Miller is performed for cell growth. Then, the cells are centrifuged, and the pellet is resuspended in 50 mL of M9 minimal medium containing the carbon source used as substrate. This type of experiment with one first phase of growing followed by one second phase of production in other medium has been used to produce several valuable compounds and it was also previously optimized for the production of naringenin (Rodrigues et al. 2020; Gomes et al. 2024). The production of *p-*coumaric acid, naringenin, 8-PN, 3’-PN, and 6-PN by the 12 producing strains was evaluated (**Tab. 2**).

**Tab. 2.** *De novo* production of PN compounds in shake flask experiments using the combination of LB Miller and M9 minimal media. Production experiments were performed in triplicate for the 12 identified strains expressing the pRSFDuet_*Fj*TAL_*Cm*CHS_*At*4CL_*Ms*CHI (NAR plasmid) and the pCDFDuet vectors holding prenyltransferases (PTs) able to de novo produce PN in the 96-well deep well plates experiments. The production of *p-*coumaric acid (CA), naringenin (NAR), 3’-prenylnaringenin (3’-PN), and 6-prenylnaringenin (6-PN) was evaluated by ultra-high performance liquid chromatography (UHPLC).

| Strain | PT | Produced compound (μM) |  |  |  |
| --- | --- | --- | --- | --- | --- |
|  |  | CA | NAR | 3'-PN | 6-PN |
| *E. coli* M-PAR-121:*BsDXS* | CdpC3PT | 219.13 ± 49.73 | 15.23 ± 1.41 | 1.54 ± 0.64 | - |
|  | CoAnaPT | 340.09 ± 32.82 | 22.94 ± 6.09 | 3.38 ± 1.43 | - |
|  | CloQ | 165.93 ± 23.26 | 6.29 ± 4.72 | 0.56 ± 0.12 | - |
|  | NphB | 1660.32 ± 84.39 | 32.10 ± 9.35 | 4.26 ± 9.35 | - |
|  | *Sp*PT | 124.44 ± 13.98 | 25.27 ± 3.57 | 29.85 ± 4.92 | - |
|  | UbiA | 58.97 ± 5.04 | 16.53 ± 3.41 | 12.45 ± 2.16 | - |
| *E. coli* M-PAR-121:*EcDXS* | CoAnaPT | 384.38 ± 42.00 | 68.90 ± 5.98 | 39.10 ± 4.35 |  |
|  | CloQ | 442.74 ± 70.39 | 137.81 ± 12.36 | 71.71 ± 7.18 | 1.66 ± 0.17 |
|  | NphB | 436.03 ± 89.08 | 11.90 ± 1.27 | 4.53 ± 0.93 | - |
|  | *Sp*PT | 219.18 ± 16.90 | 40.39 ± 9.15 | 21.90 ± 3.79 | - |
| *E. coli* M-PAR-121:*BsDXS*:*ScIDI* | CloQ | 106.48 ± 2.59 | 1.10 ± 0.48 | 2.68 ± 0.58 | - |
| *E. coli* M-PAR-121:*EcDXS*:*EcIDI* | *Ec*PT | 183.19 ± 32.67 | 23.88 ± 5.42 | 9.96 ± 1.32 | - |

PN compounds were detected for all the 12 combinations in the shake flask experiments using the combination of LB Miller and M9 minimal media. The production of 3’-PN was lower than the production achieved in 96-well deep well plates experiment for some of the combinations. As occurred in 96-well deep well plates experiments, *E. coli* M-PAR-121:*Ec*DXS expressing pRSFDuet_*Fj*TAL_*Cm*CHS_*At*4CL_*Ms*CHI and pCDFDuet_CloQ was the best producing strain being able to produce 71.71 ± 7.18 µM of 3’-PN and 1.66 ± 0.17 µM of 6-PN. Taking these results into account, this strain was chosen to proceed with further optimizations.

Due to the relevance of these compounds and the importance of implementing an industrial process, several optimizations must be carried out. One of the problems with using the LB+M9 combination in these shake flask experiments is the fact that this is not a one-step process and there is a need to recover the cells and resuspend them in the production medium. This makes the experiments more prone to contamination and is difficult to implement on a larger production scale. Moreover, the production step in M9 minimal medium does not allow the evaluation of cell growth due to the presence of CaCO_3_ in suspension. Implementing a one-step production strategy would make the production process more economically viable and easier to scale-up (Couto et al., 2017). Considering that, shake flask production experiments using the *E. coli* M-PAR-121:*Ec*DXS strain expressing pRSFDuet_*Fj*TAL_*Cm*CHS_*At*4CL_*Ms*CHI and pCDFDuet_CloQ were performed in TB and M9 modified media (**Fig. 2**). M9 modified medium differs from the M9 minimal medium used in the LB+M9 experiments due to the presence of trace elements and yeast extract instead of vitamins. Yeast extract was supplemented to improve *E. coli* growth and to increase the protein expression and production (Chen et al., 2024, 2022; Tachibana et al., 2021). TB medium was prepared as used in the 96-well deep well plates experiment.

**Fig. 2.**
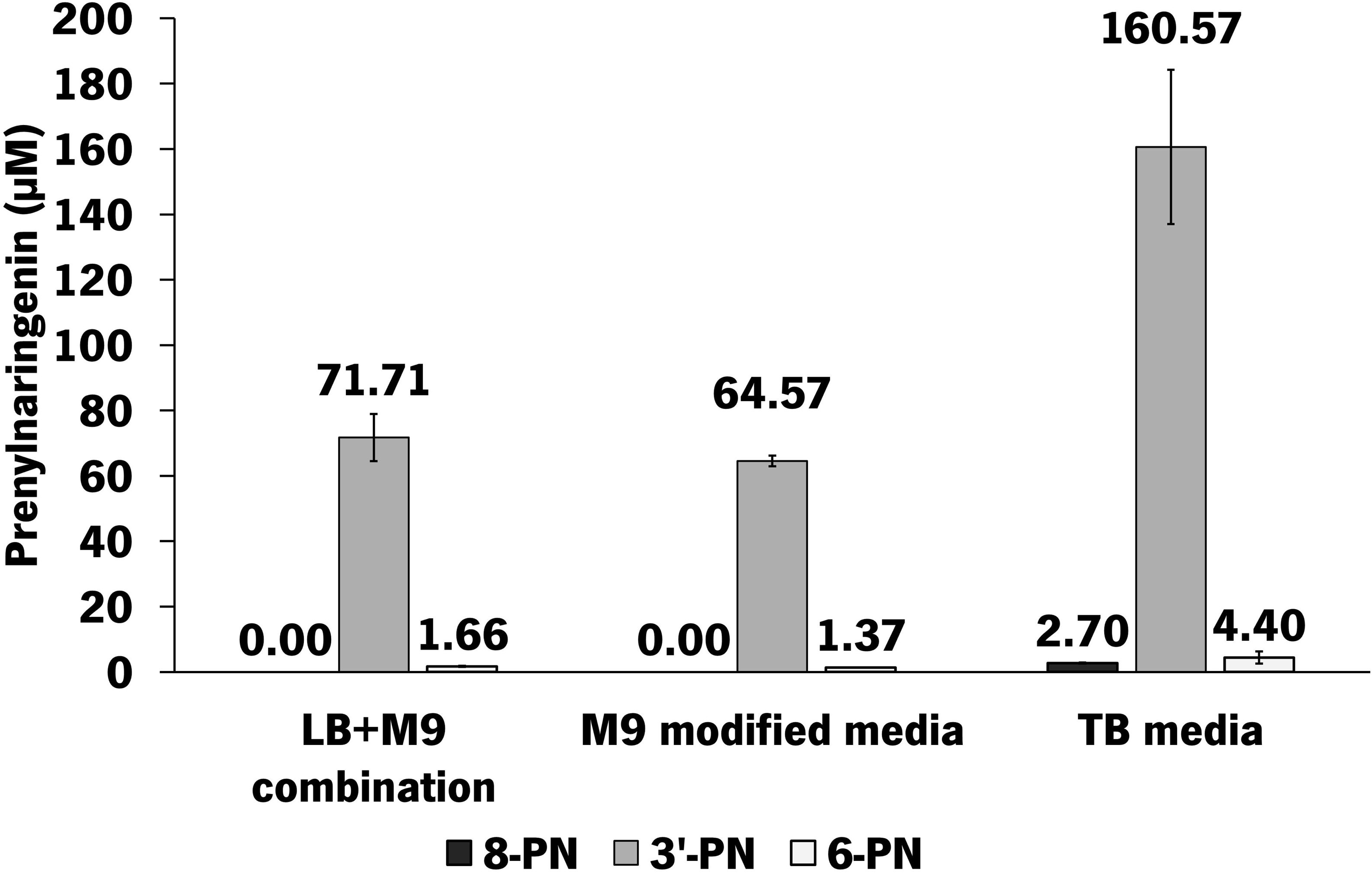

As can be observed in **Fig. 2**, the production of PN compounds was higher using TB medium. Using this production medium, *E. coli* M-PAR-121:*Ec*DXS expressing pRSFDuet_*Fj*TAL_*Cm*CHS_*At*4CL_*Ms*CHI and pCDFDuet_CloQ was able to produce 160.57 ± 23.60 µM of 3’-PN, 4.40 ± 1.85 µM of 6-PN, and 2.70 ± 0.15 µM of 8-PN. Moreover, 3’-PN is the most produced compound in 96-well deep well plates experiments and also in shake flask experiments. Nevertheless, the production of this compound was significantly improved (2.2-fold) using TB medium instead of the combination of LB+M9. In contrast, the production levels using M9 modified medium were slightly lower than the ones obtained using the combination of LB+M9. However, the differences were not statistically significant. Despite this, from an industrial and scale-up point of view, it would be more advantageous to use the M9 modified medium than the combination of LB+M9.

The production profile of all the compounds and also the profile of glucose consumption was evaluated during all the production experiments for all the production media tested (**Fig. 3**).

**Fig. 3.**
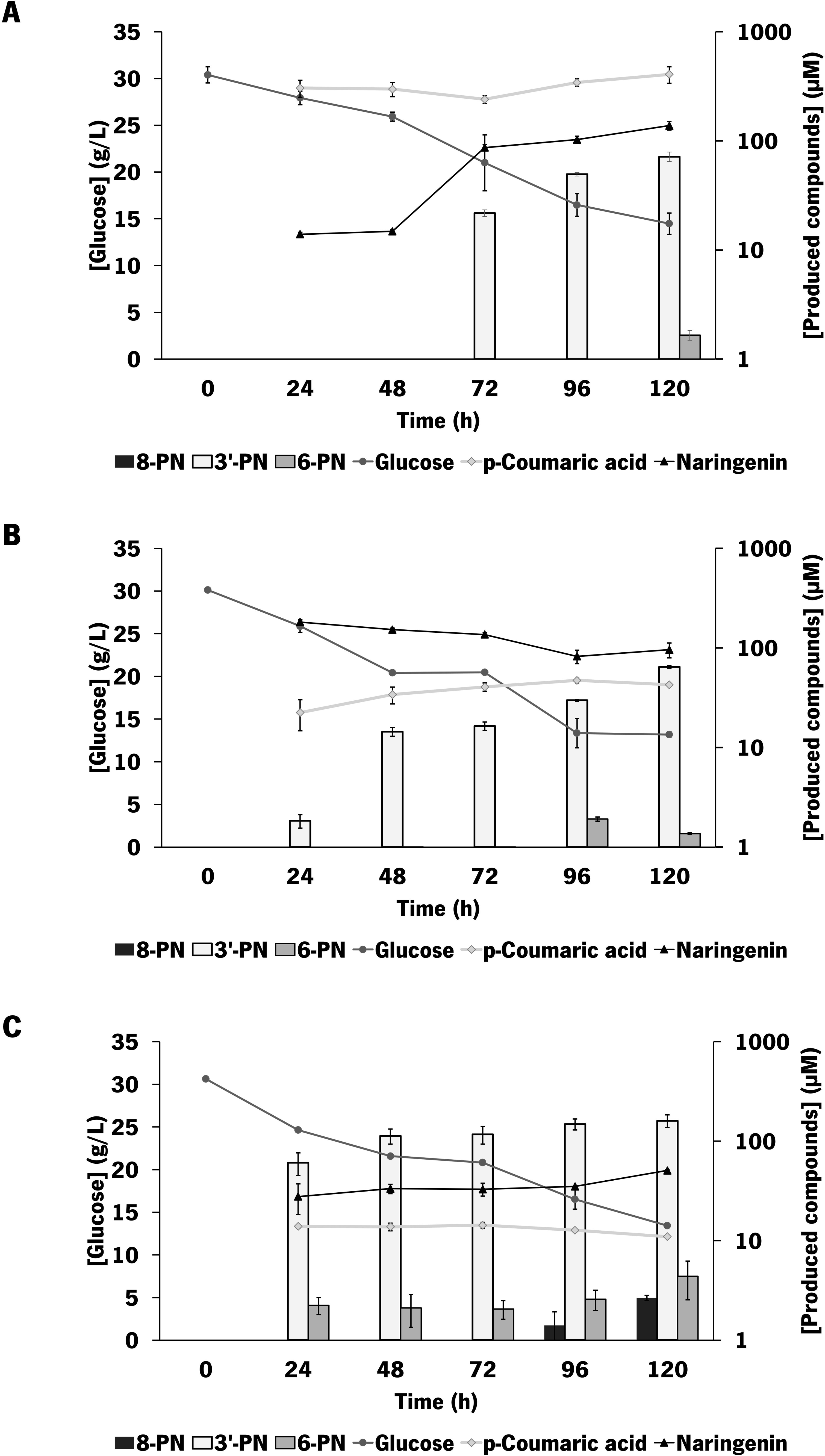

As can be observed in **Fig. 3**., the glucose consumption was similar in the tested production media. Only 16-17 g/L of glucose were consumed in these experiments. Regarding the intermediaries *p-*coumaric acid and naringenin, higher amounts of these compounds were accumulated in the production experiment using LB+M9. By other side, 3’-PN was only detected 72 h after the beginning of this experiment and 6-PN is only detected in the final time point (120 h). In the experiment using M9 modified medium, 3’-PN was detected at 24 h and the production of this compound was exponentially increasing during the experiment. As occurred in LB+M9, 6-PN was only detected in the final time points of the experiment (96 h and 120 h). Regarding TB experiment, higher amounts of 3’-PN were detected since the 24 h of the experiment and the production also increased over time. Contrarily to the other experiments, 6-PN was also detected since the 24 h of the experiment. Moreover, 8-PN was also detected in the final time points of the experiment (96 h and 120 h), being only detected in this experiment. Considering these results, the use of M9 modified or TB media can be considered advantageous compared to the use of the combination of LB+M9 since PN compounds are produced and detected earlier in the experiment.

### 3.4. Evaluation of *de novo* PN production by *Escherichia coli* M-PAR-121:EcDXS expressing pRSFDuet_*Fj*TAL_*Cm*CHS_*At*4CL_*Ms*CHI and pCDFDuet_CloQ at a bioreactor scale

With the aim of increasing production levels, the scale-up of the production process for 2-L lab-scale stirring bioreactor was considered. Since the production of 3’-PN and 6-PN was significantly higher when TB was used in shake flask experiments, this production medium was selected for bioreactor experiments. The production profile of all the compounds and glucose consumption profile were evaluated throughout the production experiment. Moreover, the cultures growth was also evaluated during the experiment (**Fig. 4**).

**Fig. 4.**
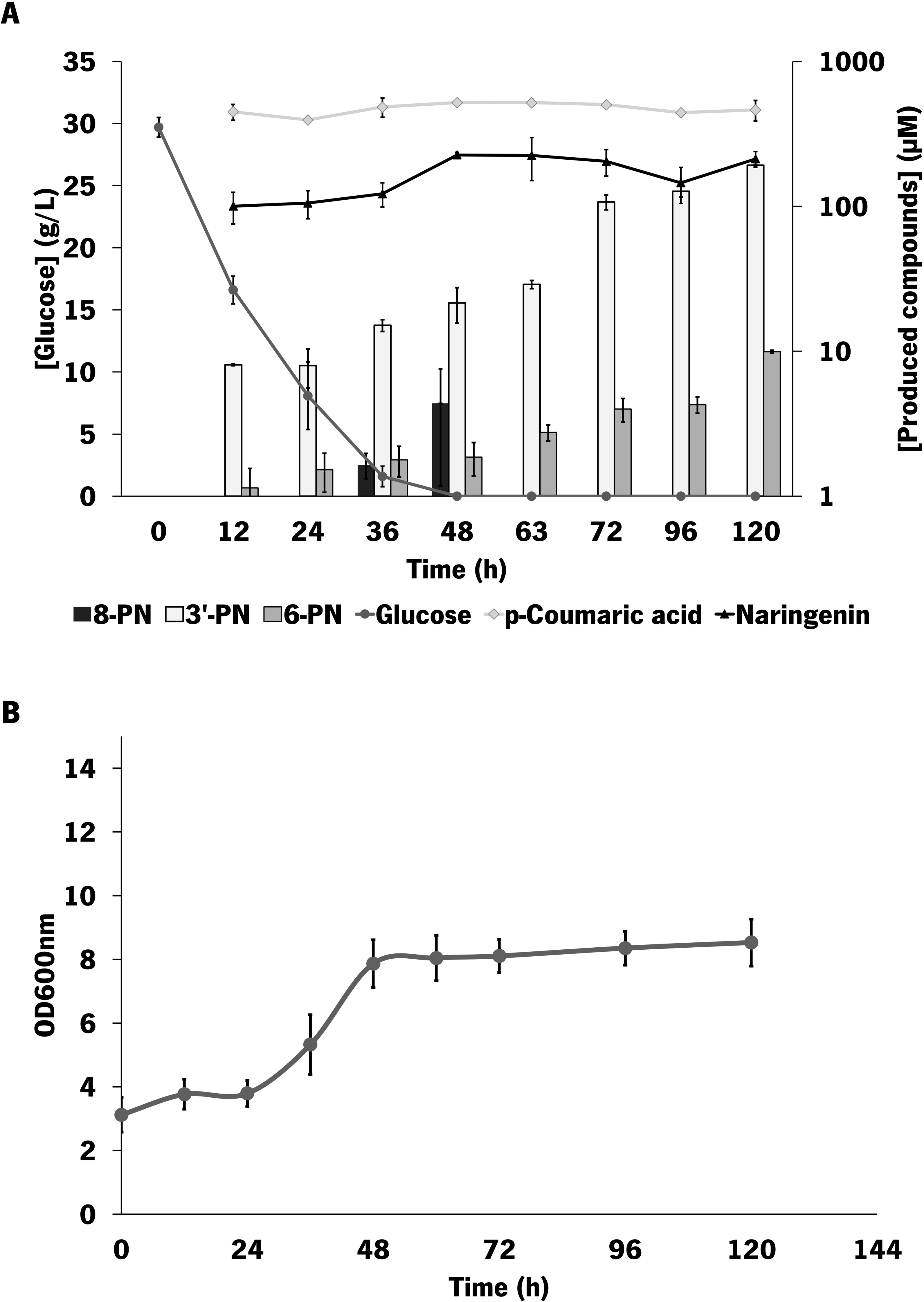

Comparing with the shake flask experiments, significantly higher amounts of the intermediates *p-*coumaric acid and naringenin were produced and accumulated during the experiment. At the final time point, 462.06 ± 74.61 µM of *p-*coumaric acid and 212.28 ± 26.54 µM of naringenin were produced, representing a 42.1-fold and 4.2-fold increase comparing with shake flask experiments, respectively. In contrast to shake flask experiments, glucose was completely consumed within the first 48 h (**Fig. 4B**). Regarding the production of PN compounds, the production of both 3’-PN and 6-PN was increasing during the experiment time and 191.94 ± 2.40 µM of 3’-PN (65.34 mg/L) and 9.9 ± 0.24 µM of 6-PN (3.37 mg/L) were detected in the final time point of the experiment. Moreover, 8-PN was detected at 36 h and 48 h. However, it was not detected in the next time points indicating that this compound was probably degraded. Although the differences in the 3’-PN production were not statistically significant compared to the shake-flask experiments, higher amounts of this compound were detected, with a 1.2-fold improvement in the production levels. Regarding 6-PN, the production achieved in the bioreactor experiment represents a 2.3-fold improvement compared to the production in shake flask experiments. However, the differences were not considered statistically significant since *p*-value is above 0.05.

Since glucose (30 g/L) was fully consumed within the first 48 h of this experiment, the addition of two extra glucose pulses (10 g/L in each pulse) was considered (**Fig. 5**). Due to the additional glucose, the experiment was extended to 144 h instead of 120 h.

**Fig. 5.**
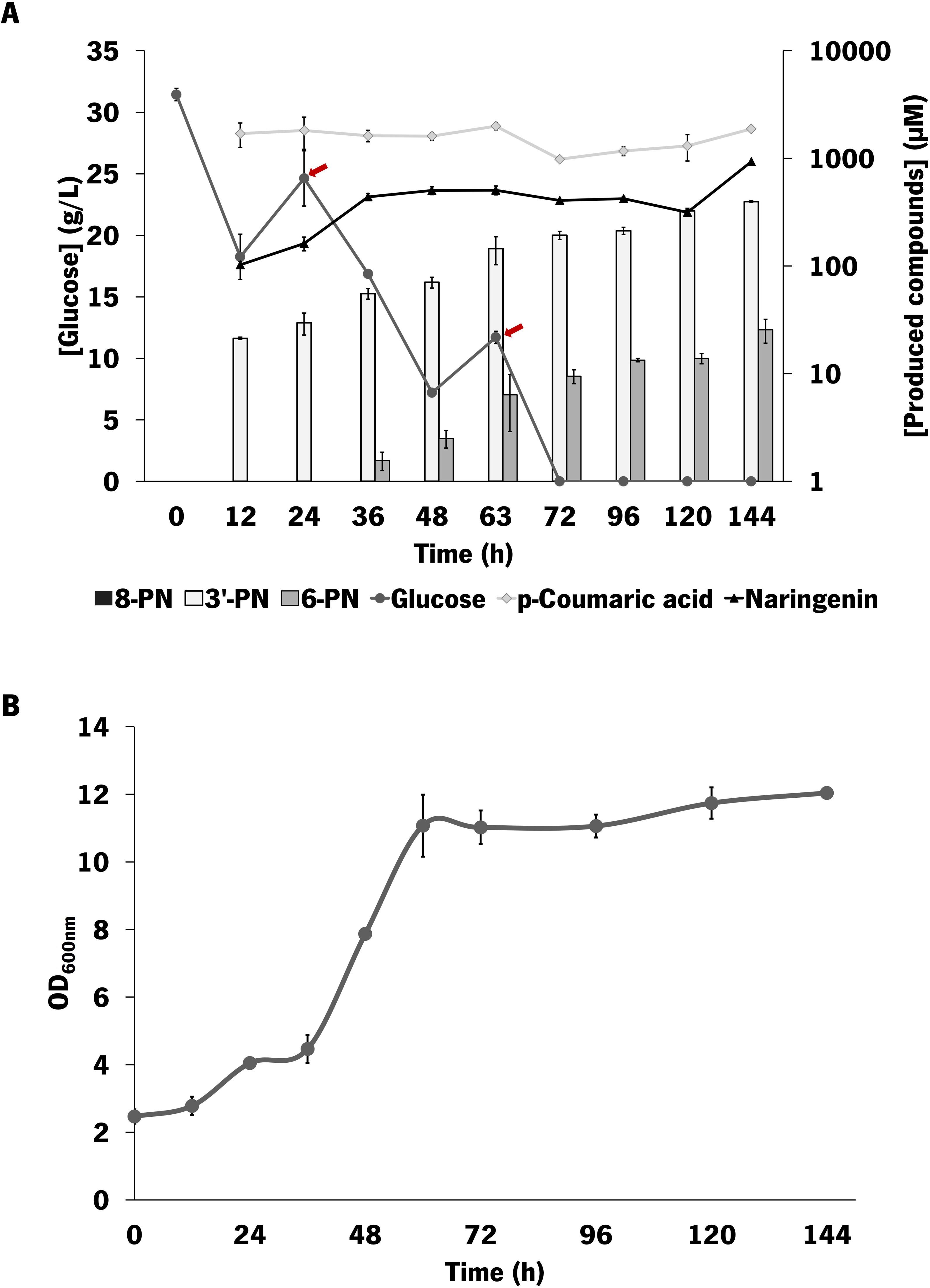

Despite the two additional glucose pulses (10 g/L), all the supplied glucose was consumed in 72 h. Compared to the shake-flask experiments and to the first batch experiment, significantly higher amounts of the intermediates *p-*coumaric acid and naringenin were produced and accumulated during the experiment. At the final time point, 1879.18 ± 73.79 µM of *p-*coumaric acid and 932.84 ± 357.07 µM of naringenin were produced. In this experiment, 8-PN was not detected in any time point. This result indicates that the compound may not be produced or may be produced in such low amounts that it is not detected or is being degraded. As observed in the first experiment, the production of both 3’-PN and 6-PN was also increasing during the experiment time. At 144 h, 397.57 ± 8.25 µM of 3’-PN (135.33 mg/L) and 25.61 ± 6.36 µM µM of 6-PN (8.72 mg/L) were produced. A representative UHPLC chromatogram of this sample and reference standard can be observed in **Fig. S7**. These results represent a 2.1-fold increase in the production of 3’-PN and a 2.6-fold increase of 6-PN compared with the first batch experiment in bioreactor, indicating that the presence of higher amounts of glucose is useful to improve the production levels.

## 4. Discussion

PN compounds have been reported as powerful compounds due to their interesting biological activities, namely anticancer and estrogenic. In this work, we aimed to design, construct and validate an *E. coli* strain able to produce PN compounds from a simple carbon source (glucose).

Since the step catalysed by the PT is considered the critical step, several PTs were selected and tested to validate their ability to perform the prenylation step. Eleven PTs were selected: three from plant sources and eight from microbial sources. Their expression was individually tested SDS-PAGE gel and as expected, only microbial aromatic PTs presented the desired protein bands (**Fig. S1 and Fig. S2**). Compared with plant aromatic PTs, microbial PTs are more easily expressed in *E. coli* due to their solubility. Despite that, all the PTs were tested towards the production of PN. Before this work, *E. coli* was only used as chassis to express PTs and to directly convert flavonoids into prenylflavonoids (Qiu et al. 2021; Liu et al. 2023; Zhang et al. 2024; Fan et al. 2025). In the bioconversion studies previously performed in *E. coli*, DMAPP availability was improved by introducing the isopentenol utilization pathway, composed by a promiscuous kinase (PK) and isopentenyl phosphate kinase (IPK). Moreover, only PTs from microbial sources were expressed. In the first study, Qiu et al. (2021) have reported the bioconversion of 200 mg/L of naringenin into 69.5 mg/L of 6-PN by expressing a PT gene from *Streptomyces sp.* NT11 (*Sh*FPT) in combination with the PK from *Shigella flexneri* (*Sf*PK) and IPK from *Methanolobus tindarius* (*Mt*IPK) to improve DMAPP flux. Later, Liu et al. (2023) reported the bioconversion of 200 mg/L of naringenin into 10.3 mg/L of 3’-PN. This result was achieved by expressing a bacterial PT from *Nocardiopsis gilva* (*Ng*FPT) in combination with *Sf*PK and *Mt*IPK. Moreover, Zhang et al (2024) reported higher bioconversion levels using the AnaPT from *N. fischeri* to catalyse the prenylation step and PK from *E. coli* (*Ec*PK) and IPK from *A. thaliana* (*At*IPK) to improve DMAPP flux. In a first approach, shake flask experiments were performed using 300 mg/L of naringenin as substrate and 142.10 mg/L of 3’-PN were produced. Moreover, a fed-batch strategy was tested at a bioreactor scale and 537.80 mg/L of 3’-PN with the feeding of 200 mg/L naringenin in three time points (6 h, 18 h and 36 h). More recently, *E. coli* was also engineered towards the bioconversion of other flavonoids skeletons into their prenylated forms. Several PTs from fungal species were tested and Ad03 from *Aspergillus nidulans* and Ao01 from *Aspergillus oryzae* were selected due to their prenylation ability. DMAPP availability was also improved by introducing *Ec*PK and IPK from *Thermoplasma acidophilum* (*Ta*IPK). When 200 mg/L of silybin were supplemented, the engineered strain expressing Ad03 was able to produce 176 mg/L of 6-prenylsilybin. When 200 mg/L of daidzein were supplemented, the same strain was able to produce 53.5 mg/L of the prenylated compound erysubin F. Moreover, the engineered strain expressing Ao01 was able to convert 200 mg/L of baicalein into 32 mg/L of 6-*O*-prenylbaicalein (Fan et al., 2025). However, the supplementation of naringenin or other flavonoids in these bioconversion studies increases the production costs associated with the process. At an industrial scale, this supplementation will lead to an economically unviable process. For this reason, assembling the complete biosynthetic pathway to produce PN compounds from a simple and cheap carbon source, such as glucose, is mandatory to achieve a cost-effective production process. Considering this, *E. coli* M-PAR-121 strains carrying pRSFDuet_*Fj*TAL_*Cm*CHS_*At*4CL_*Ms*CHI and pCDFDuet_PT vectors were tested to evaluate PN production from glucose. However, no production of any PN compound was detected in any combination possibly due to a limitation in the DMAPP availability. The improvement of DMAPP availability was considered to try to achieve the production of PN from glucose. As previously mentioned, the isopentenol utilization pathway was engineered to improve DMAPP in the bioconversion studies. However, the utilization of this pathway requires the exogenous supplementation of prenol. This supplementation increases the production costs, limiting the further application and scale-up to an industrial scale (Liu et al., 2023; Qiu et al., 2021; Zhang et al., 2024). Taking this into account, in this study, we have focused on the improvement of DMAPP supply by eliminating the previously identified rate-limiting steps DXS and IDI of the MEP pathway responsible for DMAPP synthesis in *E. coli*, using CRISPR-Cas12a (Banerjee et al., 2013; Li et al., 2017; Rinaldi et al., 2022; Yuan et al., 2006; Zhou et al., 2012). This strategy has the advantage of not needing the supplementation of any precursor. Single and double integration of DXS and IDI genes with improved activities (*Bs*DXS, *Sc*IDI, and *Bl*IDI) were performed into *E. coli* M-PAR-121 genome. The overexpression of these genes was previously tested and proved to be useful to eliminate the bottleneck steps of the MEP pathway and resulted in the improvement of terpenoids production (Zhao et al. 2011; Rad et al. 2012; Yang and Guo 2014; Gao et al. 2016). Single and double integration of the native DXS and IDI genes from *E. coli* were also tested to increase the native flux of the pathway. In addition to improving the MEP pathway flux, the prevention of the pathway deviation can be crucial to improve the production levels. Considering that, CRISPRi was used to downregulate the *ispA* gene responsible for DMAPP conversion into GPP and FPP. Ten boosted DMAPP-*E. coli* strains were successfully constructed and used as chassis for the expression of the complete PN pathway. *De novo* production of PN was firstly screened in 96-well deep well plates since it enables a fast screening of several strains in simultaneous. As can be observed in **Tab. 3**, *de novo* production of PN was not detected in the wild-type strain *E. coli* M-PAR-121, corroborating the expected DMAPP limitation. Moreover, the strains holding only the integration of IDI gene were not able to *de novo* produce PN compounds indicating that the overexpression of the last step gene of the MEP pathway may not be enough to increase the flux for DMAPP production. In fact, it is reported that the DXS step is the most regulated step in the MEP pathway and that the overexpression of this gene can result in the elimination of this bottleneck and in the improvement of the flux towards DMAPP synthesis (Banerjee et al., 2016; Gao et al., 2016; Rinaldi et al., 2022). In addition to these strains, the two strains constructed with CRISPRi technology with different levels of downregulation of *ispA* gene were also not able to produce PN compounds. This probably occurred due to a high metabolic burden on the cells since three plasmids are being simultaneously expressed. In the future, it will be useful to integrate also the CRISPRi machinery present on the CRISPRi plasmid on the *E. coli* genome to reduce the cells metabolic burden and stably maintain the repression levels, removing also the need of another antibiotic supplementation. Moreover, the repression of an essential gene can be affecting the growth and the metabolic balance of *E. coli* cells (Jervis et al., 2021). In fact, it was verified in the qPCR experiments that *E. coli* M-PAR-121 with pBbS8c-*ddcpf1*-*ispA*1 and *E. coli* M-PAR-121 with pBbS8c-*ddcpf1*-*ispA*2 had lower growth rates compared to the control strain, corroborating that *ispA* repression was affecting the cellular growth (*data not shown*).

Regarding the eleven PTs tested in this study, *de novo* production of PN compounds was not detected when any of the plant aromatic PTs were expressed probably due to the lower solubility of these enzymes and the presence of signal peptides that target them to the biological membranes. This result was already expected since *E. coli* does not have intracellular compartments that are usually required for the efficient expression of these plant PTs (de Bruijn et al., 2020). Contrary to plant aromatic PTs, microbial PTs are soluble enzymes being more easily expressed in *E. coli.* All the microbial PTs apart from AnaPT were able to produce PN compounds. However, PN was detected when the codon-optimized version of this enzyme was also tested (coAnaPT), showing the importance of codon optimization to enable the efficient expression of the target proteins (Rainha et al., 2020; Rodrigues and Rodrigues, 2017). Shake flask experiments using a combination of LB+M9 were also performed for the 12 identified producing strains (**Tab. 4**). *E. coli* M-PAR-121:*Ec*DXS expressing pRSFDuet_*Fj*TAL_*Cm*CHS_*At*4CL_*Ms*CHI and pCDFDuet_CloQ was identified as the best producing strain in both experiments (96-well deep well plate and shake flask experiments). Previous studies have reported the successful expression of CloQ, the aromatic PT from *S. roseochromogenes*, in *E. coli* and its utilization for clorobiocin production (Araya-Cloutier et al., 2017; Metzger et al., 2010; Pojer et al., 2003). Among the PN compounds analysed (8-PN, 3’-PN, and 6-PN), higher production of 3’-PN was achieved. In fact, CloQ, was previously characterized and it was reported to perform mostly C-prenylation in the B ring of the flavonoid skeleton using DMAPP as donor substrate (Araya-Cloutier et al., 2017). This PT is also able to perform prenylations in the A ring but with lower regioselectivity. This preference for the C-prenylation in the B ring can be the justification for the higher productions of 3’-PN (Araya-Cloutier et al., 2017). Moreover, contrarily to most of the PTs, CloQ was found to be independent of the presence of metallic ions, namely Mg^2+^. This characteristic can be useful and advantageous because it is one less factor affecting PT activity (Metzger et al., 2010). 3’-PN and 6-PN production attained in 96-well deep well plates using TB medium was slightly higher than the production achieved in the shake flask experiments using LB+M9. This result could indicate that TB medium favours the PN production probably due to a higher cell growth in this medium. Moreover, the use of the LB+M9 combination is not feasible for the implementation of the production process at an industrial level due to the need to exchange the medium during the experiment. Considering that, shake flask experiments were performed in TB and M9 modified media for the best producing strain. Higher productions were achieved in TB corroborating that this medium is better to efficiently produce PN compounds (**Fig. 2** **and** **Fig. 3**). This result can be justified by the higher *E. coli* growth rates reached in TB medium comparing with the obtained in M9 modified medium (OD_600nm_=2.9 in TB medium vs OD_600nm_=2.1 in M9 modified medium (*data not shown*)). Then, this medium was selected to scale-up the production process in a 2-L lab-scale stirring bioreactor. In the initial batch experiment, higher amounts of 3’-PN and 6-PN were produced comparing with the shake flask experiments (**Fig. 4**). The production of the intermediaries *p-*coumaric acid and naringenin was also higher. At the final time point, 191.94 ± 2.40 µM of 3’-PN (65.34 mg/L) and 9.9 µM of 6-PN (3.37 mg/L) were produced. These results can be justified by the significantly higher growth attained in the bioreactor (OD_600nm_=8.5 in batch experiment in bioreactor vs OD_600nm_ =2.9 in shake flask experiments). Comparing with shake flasks, bioreactors provide a highly controlled environment allowing the control of several parameters that are essential for optimal *E. coli* growth, namely pH, dissolved oxygen and agitation. For example, the oxygen transfer in shake flasks can be more limiting due to inconsistent mixing. Moreover, the bioreactor system used in this study has sensors to control key parameters in real-time, namely pH and dissolved oxygen, being automatically adjusted for the setting parameters. This represents an advantage comparing with shake-flasks because the optimal conditions for *E. coli* growth are constantly maintained.

The increase in the growth rate in the bioreactor experiment was accompanied by a full glucose consumption in the first 48 h of the experiment. Considering the full glucose consumption, another bioreactor experiment was performed providing two additional glucose pulses (10 g/L) (**Fig. 5**). Higher amounts of all the metabolites were produced and accumulated comparing with the first experiment. In fact, the two additional glucose pulses resulted in an improvement in the cellular growth to a final OD_600nm_ of 12.0, being the main reason for the higher production levels that were achieved. Regarding PN compounds, 397.57 ± 8.25 µM of 3’-PN (135.33 mg/L) and 25.61 ± 6.36 µM µM of 6-PN (8.72 mg/L) were produced. As far as we know, until this study, only *S. cerevisiae* was engineered towards *de novo* production of PN compounds. *De novo* production of 8-PN was achieved by Levisson et al. (2019) and Guo et al. (2022). Firstly, Levisson et al. (2019) have integrated the complete naringenin biosynthetic pathway derived from *A. thaliana* into the genome of a *L-*phenylalanine and *L-*tyrosine *S. cerevisiae* overproducing strain. Then, this strain was used to express the flavonoid PT from *S. flavescens* (SfPT). DMAPP flux was improved by expressing a truncated variant of the HMGR (tHMG1), that converts 3-hydroxy-3-methylglutaryl-CoA (HMG-CoA) to MVA. Moreover, enoyl-CoA reductase (ECR) from *Malus domestica* was used to replaced the native TSC13 to prevent the deviation of *p*-coumaroyl-CoA (intermediary of naringenin biosynthetic pathway) into phloretic acid. This strain was able to produce 0.12 mg/L of 8-PN using 20 g/L of glucose as substrate. Higher 8-PN production levels were further achieved by Guo et al. (2022). In this study, the authors have performed enzyme engineering to improve the efficiency of naringenin 8-dimethylallyltransferase 1 (N8DT-1) from *S. flavescens* (*Sf*N8DT-1). In a first approach, this enzyme was truncated to remove signal peptides. Then, key residues involved in the catalytic efficiency were identified and mutated (Q12E and N305M). Moreover, the DMAPP flux was improved by expressing tHMGR and IDI. The combination of all these strategies resulted in the production of 49.35 mg/L and 101.40 mg/L of 8-PN from glucose in shake flask experiments and 5 L bioreactor, respectively. This 8-PN production is the highest reported so far for this compound being in the same range of production achieved in our study for 3’-PN.

Regarding 3’-PN, this compound was only produced in *S. cerevisiae* using *L-*phenylalanine as substrate (Isogai et al., 2021). In a first approach, codon-optimized *Sf*N8DT-1 and the acetyl-CoA carboxylase (ACC) gene (to improve malonyl-CoA) were expressed in a yeast platform strain holding in the genome the naringenin biosynthetic pathway derived from *A. thaliana.* However, only 0.615 μg/L of 3’-PN were produced from *L-*phenylalanine. Moreover, alternative aromatic PTs were also expressed and 1.10 μg/L of 3’-PN were obtained when AnaPT was expressed (Isogai et al., 2021). This production levels using *L-*phenylalanine as substrate are far away from the production obtained in our study.

As far as we know, our work represents the first report of *de novo* production of PN compounds using *E. coli* as chassis without any supplementation of the intermediate naringenin and the first study of *de novo* production of 3’-PN in any microbial chassis. Despite that, the production levels herein reported are still far from what is needed to satisfy industrial demand. Taking this into account, further optimizations should be considered. For example, other strategies to improve DMAPP availability, such as the reconstruction of a heterologous MVA pathway and other strategies of downregulation of competing pathways that deviate DMAPP should be implemented (Cao et al., 2019). Moreover, PT step should also be optimized since this is the most critical step of the pathway. Signal sequences and relevant amino acids that are involved in catalytical activity and that interact with donor and acceptor substrates should be investigated and strategies of rational design, directed evolution and random or directed mutagenesis should be considered to improve the catalytical efficiency of the PT enzyme and, consequently, the production levels (Isogai et al., 2021; Ko et al., 2020; Wu et al., 2020).

## CRediT authorship contribution statement

DG: Investigation; Methodology; Conceptualization; Writing - Original Draft; Writing - Review and Editing; Visualization. JR: Methodology; Conceptualization; Validation; Writing - Review and Editing; Supervision. NS: Resources; Funding acquisition; Writing - Review and Editing; LR: Conceptualization; Resources; Funding acquisition; Writing - Review and Editing; Supervision.

## Acknowledgements

The authors thank Dr. Daisuke Koma for kindly providing the *E. coli* M-PAR-121 strain (Koma et al. 2020). The authors also thank Dr. Shu-Ming Li for providing pWY16 and pWY24 plasmids (Yin et al., 2010, 2009).

