## Supplementary material for "*De novo* production of prenylnaringenin compounds by a metabolically engineered *Escherichia coli*"

**Table 1. Strains used in this study.**

| **Strains** | **Relevant Genotype** | **Source** |
| --- | --- | --- |
| *E. coli* NZY5α | fhuA2Δ(argF^-^lacZ)U169 phoA glnV44 Φ80 Δ(lacZ)M15 gyrA96 recA1 relA1 endA1 thi-1 hsdR17 | NZYTech (MB00401) |
| *E. coli* NEB5α | fhuA2Δ(argF-lacZ)U169 phoA glnV44 Φ80Δ(lacZ)M15 gyrA96 recA1 relA1 endA1 thi-1 hsdR17 | (New England Biolabs - C2987H) |
| *E. coli* M-PAR-121 | *L-*Tyrosine overproducing strain derived from MG1655 (DE3); *tyrR*::P_T7lac_-*aroG^fbr^ ldhA*::P_T7lac_-*tyrA^fbr^ adhE*::P_T7lac_-*ppsA pflDC*::P_T7lac(-8TC)_-*tktA pykF*::P_T7lac_-*aroALC ascF*::P_T7lac_-*aroEDB* | (Koma et al., 2020) |
| *E. coli* M-PAR-121:*Bs*DXS | *E. coli* M-PAR-121 with the integration of 1-deoxy-D-xylulose-5-phosphate synthase (DXS) gene from *Bacillus subtilis* (*Bs*DXS) into the *lacZ* locus of the genome | This study |
| *E. coli* M-PAR-121:*Sc*IDI | *E. coli* M-PAR-121 with the integration of isopentenyl diphosphate isomerase (IDI) from *Saccharomyces cerevisiae* (*Sc*IDI) into the *lacZ* locus of the genome | This study |
| *E. coli* M-PAR-121:*Bl*DI | *E. coli* M-PAR-121 with the integration of IDI from *Bacillus licheniformis* (*Bl*IDI) into the *lacZ* locus of the genome | This study |
| *E. coli* M-PAR-121:*Bs*DXS-*Sc*IDI | *E. coli* M-PAR-121 with the integration of *Bs*DXS and *Sc*IDI into the *lacZ* locus of the genome | This study |
| *E. coli* M-PAR-121:*Bs*DXS-*Bl*DI | *E. coli* M-PAR-121 with the integration of *Bs*DXS and *Bl*IDI into the *lacZ* locus of the genome | This study |
| *E. coli* M-PAR-121:*Ec*DXS | *E. coli* M-PAR-121 with the integration of DXS gene from *Escherichia coli* (*Ec*DXS) into the *lacZ* locus of the genome | This study |
| *E. coli* M-PAR-121:*Ec*IDI | *E. coli* M-PAR-121 with the integration of IDI gene from *E. coli* (*Ec*IDI) into the *lacZ* locus of the genome | This study |
| *E. coli* M-PAR-121:*Ec*DXS-*Ec*IDI | *E. coli* M-PAR-121 with the integration of *Ec*DXS and *Ec*IDI into the *lacZ* locus of the genome | This study |

Table 2. Plasmids used in this study.

| **Plasmids** | **Construct** | **Source** |
| --- | --- | --- |
| pCDFDuet-1 | CloDF13 ori, lacI, double PT7lac, Spec^R^ | Novagen |
| pRSFDuet_*Fj*TAL_*Cm*CHS | pRSFDuet-1 (RSF1030 ori, lacI, double PT7lac, Kan^R^) carrying codon-optimized tyrosine-ammonia lyase (TAL) from *Flavobacterium johnsoniae (Fj*TAL*)* and chalcone synthase (CHS) from *Curcubita maxima* (*Cm*CHS) | (Gomes et al. 2024) |
| pACYCDuet_*At*4CL_*Ms*CHI | pACYCDuet-1 (P15A ori, lacI, double PT7lac, Cm^R^) carrying 4-coumarate:CoA ligase 1 (4CL-1) from *Arabidopsis thaliana* (*At*4CL) and chalcone isomerase (CHI) from *Medicago sativa* (*Ms*CHI) | (Gomes et al. 2024) |
| pWY16 | pGEM-T easy vector (F1 ori, Cm^R^) carrying the prenylatransferase (PT) AnaPT from *Neosartorya fischeri* | (Yin et al., 2009) |
| pWY24 | pGEM-T easy vector (F1 ori, Cm^R^) carrying CdpC3PT from *N. fischeri* | (Yin et al., 2010) |
| SBC104673 | pBbe5k vector (p15A ori, Kan^R^) carrying a codon-optimized version of PT3 from *Cannabis sativa* (*Cs*PT3) | Scrutton laboratory. |
| SBC104676 | pBbe5k vector (p15A ori, Kan^R^) carrying a codon-optimized version of AnaPT (coAnaPT) | Scrutton laboratory. |
| SBC104690 | pBbe5k vector (p15A ori, Kan^R^) carrying a codon-optimized version of CloQ from *Streptomyces roseochromogenes* | Scrutton laboratory. |
| SBC104687 | pBbe5k vector (p15A ori, Kan^R^) carrying a codon-optimized version of PT from *E. coli (Ec*PT*)* | Scrutton laboratory. |
| SBC104679 | pBbe5k vector (p15A ori, Kan^R^) carrying a codon-optimized version of NphB from *Streptomyces* sp. | Scrutton laboratory |
| SBC104684 | pBbe5k vector (p15A ori, Kan^R^) carrying a codon-optimized version of PT from *Streptomyces sp.* Act143 (*Sp*PT) | Scrutton laboratory |
| SBC104693 | pBbe5k vector (p15A ori, Kan^R^) carrying a codon-optimized version of UbiA from *E. coli* | Scrutton laboratory |
| pSIM*cpf1* | pSIM18 derived vector, Rep101 ori, HygR, P_BAD_-α-pMB1 array, P_JS23151_ *Ascpf1* | (Jervis et al., 2021) |
| pTF-lacZ-rfp | pMB1 ori, Spec^R^, P_JS23119_-*lacZ* array, *lac*Z::*rfp* | (Jervis et al., 2021) |
| pBbS8c-*ddcpf1*-∆ | SC101 ori, Cm^R^, PBAD-*ddcpf1*, P_JS23119_-empty | (Jervis et al., 2021) |
| pRSFDuet_*Fj*TAL_*Cm*CHS_ *At*4CL_*Ms*CHI | pRSFDuet-1 (RSF1030 ori, lacI, double PT7lac, Kan^R^) carrying *Fj*TAL*, Cm*CHS*, At*4CL, and *Ms*CHI | This study |
| pCDFDuet_*Hl*PT | pCDFDuet-1 carrying codon-optimized PT from *Humulus lupulus* (*Hl*PT) | This study |
| pCDFDuet_*Sf*N8DT-1 | pCDFDuet-1 carrying codon-optimized N8DT-1 from *Sophora flavescens* (*Sf*N8DT-1) | This study |
| pCDFDuet_AnaPT | pCDFDuet-1 carrying AnaPT | This study |
| pCDFDuet_CdpC3PT | pCDFDuet-1 carrying CdpC3PT | This study |
| pCDFDuet_*Cs*PT3 | pCDFDuet-1 carrying *Cs*PT3 | This study |
| pCDFDuet_CoAnaPT | pCDFDuet-1 carrying coAnaPT | This study |
| pCDFDuet_CloQ | pCDFDuet-1 carrying CloQ | This study |
| pCDFDuet_*Ec*PT | pCDFDuet-1 carrying *Ec*PT | This study |
| pCDFDuet_NphB | pCDFDuet-1 carrying NphB | This study |
| pCDFDuet_*Sp*PT | pCDFDuet-1 carrying *Sp*PT | This study |
| pCDFDuet_UbiA | pCDFDuet-1 carrying UbiA | This study |
| pTF-*lacZ*-*Bs*DXS | pMB1 ori, Spec^R^, P_JS23119_-*lacZ* array, *lac*Z::*Bs*DXS | This study |
| pTF-*lacZ*-*Sc*IDI | pMB1 ori, Spec^R^, P_JS23119_-*lacZ* array, *lac*Z::*Sc*IDI | This study |
| pTF-*lacZ*-*Bl*IDI | pMB1 ori, Spec^R^, P_JS23119_-*lacZ* array, *lac*Z::*Bl*DI | This study |
| pTF-*lacZ*-*Bs*DXS-*Sc*IDI | pMB1 ori, Spec^R^, P_JS23119_-*lacZ* array, *lac*Z::*Bs*DXS::*Sc*IDI | This study |
| pTF-*lacZ*-*Bs*DXS-*Bl*IDI | pMB1 ori, Spec^R^, P_JS23119_-*lacZ* array, *lac*Z::*Bs*DXS::*Bl*IDI | This study |
| pTF-*lacZ*-*Ec*DXS | pMB1 ori, Spec^R^, P_JS23119_-*lacZ* array, *lac*Z::*Ec*DXS | This study |
| pTF-*lacZ-Ec*IDI | pMB1 ori, Spec^R^, P_JS23119_-*lacZ* array, *lac*Z::*Ec*IDI | This study |
| pTF-*lacZ*-*Ec*DXS-*Ec*IDI | pMB1 ori, Spec^R^, P_JS23119_-*lacZ* array, *lac*Z::*Ec*DXS::*Ec*IDI | This study |
| pBbS8c-ddcpf1-ispA1 | SC101 ori, Cm^R^, PBAD-*ddcpf1*, P_JS23119_-*ispA_*array1 | This study |
| pBbS8c-ddcpf1-ispA2 | SC101 ori, Cm^R^, PBAD-*ddcpf1*, P_JS23119_-*ispA_*array2 | This study |

**Table S3. Sequence of the prenyltransferases (PTs) tested in this study.**

| **Gene** | **Sequence** |
| --- | --- |
| *Hl*PT | atggaattgtcatccgtgtcatcgttcagcctcgggaccaaccccttcattagtattcctcacaacaacaacaacttaaaagtttcctcatattgctgcaagagtaaaagtcgtgttatcaacagcaccaattccaaacactgcagtccgaataacaacaataatacaagcaataagacgacgcaccttctgggactttatggccaatctcgctgtttgttgaagccattgtccttcatatcctgtaatgatcagcgtgggaacagtatcagagccagcgcccagatcgaggaccgtccacccgagagcggtaacctgtcagcgttaactaacgtaaaggatttcgtgtctgtctgctgggaatacgtgcgtccctatactgctaagggtgtaatcatttgttcgtcatgcttgttcgggcgtgagctgctggaaaatcctaacctgttctcacgcccacttatcttccgggccctgctgggcatgttagccatcttaggatcatgcttctacactgccggtattaaccagatcttcgacatggacatagatcgcattaataagcctgaccttccgttagtctctggcagaatctcagttgagtctgcatggctcctgacattgagccccgccatcataggttttatcctgatactgaagctgaatagcggccctctgctgacgagcttatattgcctggctatattatcaggcaccatttacagtgtaccaccgttccgttggaagaagaacccgatcacggcgttcttatgcatactgatgatccacgcgggcctgaatttcagcgtctactacgcttcacgggccgcgctgggcttagctttcgcgtggtcccctagcttcagttttattacagcgttcataaccttcatgaccctgactctcgccagcagtaaggacctgagcgatattaacggtgaccgtaaattcggggtggagacattcgctactaaattaggggcgaagaatatcactctgctgggtacaggtctgctgttactgaattacgtggcggcgatctcaacagcaataatttggcccaaagcctttaaaagcaatatcatgttactgtcacacgccattcttgcgttctctctgatctttcaagcccgtgaactggaccgcaccaattatacacctgaggcatgtaagtcgttttacgagtttatatggattctgttttccgcagagtatgtagtgtacctctttatatga |
| *Sf*N8DT-1 | atgggttcaatgctgttagcctcatttccgggtgcaagctcaataacgaccggcggctcatgtatgcgtagcaagcagtacgctaagaactacaacgcctcgtcctacgtcactacactgtggcataagaagggaaagattcagaaggagcactgcgctgttattttctcaaaacacaacttaaagcagcactataaggtgaatgaaggtggcagtactagcaagaagcgcgaaaagaagtacacggtaaacgcgatctcggaagagagtttcgagtacgagccacaggttcgtgacccggagagtatttggggtagcgttaatgacgcactggacacgttttataaattctgtcgaccttacgctatgttctcgattgtcttaggggcaaccttcaaatcatttgtagccgtagaaaagttatcagacctcagccttacattcttcatagggtggcttcaagtcgtagtggcggttatatgcatacacatattcggagtcggactgaaccagctctgcgacatcgagatcgataaaatcaataaaccggacttaccgctggcgagtggtaagttatctttccgtaacgtagttataatcaccgccagctcactgatacttggacttgggttcgcgtggatagttgggagctggcctctgttctggacggtgctgatctgctgcatgttcactgcagcatacaacgtagacctgccgctgctgcgctggaagaagtacccagtattgacggccatcaatttcattgcggacgtagcggtgacgcgtagcttaggcttcttcctgcacatgcaaacctgcgtgtttaaacgtccgacgactttcccgcgacctttaatattctgcacagctatagtgtctatatacgccattgtcattgctttatttaaagacattcccgatatggaaggcgacgagaaattcggtattcagagcttaagcctgcgcctgggccccaagagagtcttctggatctgcgtgtccctgctggagatggcgtacggtgttacaatcctggtaggggccaccagcccgattctgtggtccaagatcataactgtgcttgggcacgcggtgttggcctcagttctttggtatcacgcaaagtcagtcgacttaacctcgaacgtagtacttcagtcattttacatgttcatctggaaattacacaccgctgactatttcttgatcccgctgttccgctaa |
| AnaPT | atgcctcccttgtctatgcaaaccgattctgttcaaggcactgctgagaataagtccctggagacgaatgggacctcaaacgaccagcaactgccgtggaaggtgctggggaaatcgcttggacttccgacaattgagcaggagcagtattggctcaacactgccccgtactttaataatcttctgatccagtgcggctacgatgtccaccagcagtaccagtacctggccttctatcaccggcatgttctccctgtccttggtccgttcattcggtccagtgcagaagccaactacatcagcggcttctcagccgagggctatcccatggaattgagtgtcaactaccaagcctccaaggcgaccgtccgactgggttgtgagcctgtgggcgagttcgcgggcacgtcgcaggatcccatgaaccagttcatgacgcgggaggttctaggaaggctctcccgtttggaccctacgtttgacctgcgcttgtttgactatttcgactcgcagtttagtttgaccacgagcgaggctaaccttgccgcgtccaagttgatcaaacagcggcggcagagcaaggtcattgcgtttgatctcaaggatggagcgatcattccaaaggcatactttttcctcaagggcaagtctcttgcgagtggaataccggtgcaggacgtcgccttcaatgctattgagtcaatcgccccgaagcaaattgaatctcccttgcgtgtcctgagaaccttcgtcaccaagctcttttcaaagcctactgtcaccagcgacgttttcattctagcggtagattgcatcgttccggaaaagtcgcgtatcaagctgtatgttgccgattcccagctgagtctggcgacgctgcgagaattctggactttgggcggatcggtcaccgactctgccacgatgaaaggactggagattgccgaagagttatggaggattctgcagtatgatgatgccgtttgttcacactcaaacatggaccagctaccgcttgtggtcaattacgagctatcgtcgggtagcgcaacgcccaagccacagctgtatcttcctttgcacggaaggaatgacgaggccatggctaatgcgctcaccaaattttgggactacttggggtggaagggattggctgcgcagtacaagaaggatctctatgcgaacaacccttgtcgaaacctcgcagagaccactacagtccagcgctgggtggccttttcttacacagagtccggtggcgcatatctgaccgtttacttccatgcagttggtggtatgaagggcaatctctagaagctt |
| CdpC3PT | atgacagtgtcgtcgaccgccgttgaggcctctgcgccctgtgctgaaatgggacatgacatcccatatcgcacactgagcaggtcaatgatattcgccaatctggatcagtatcaatactggcatcaaattggcccggtgcttggcaagatgctggtggatggggaatacagcatccaccgccaatacgagtacctgtcgctctttgctcacttgatcatccccaagctggggcccttccccagccccgggagagacatctacaggtgtctcctcggcaacattggtggtcccttcgagctcagtcagaacttccagagactcggctcaacggctcggctcgccttcgagcccacgagttatctcgccagcacgagtgccgacccattcaaccgccacgccgtccacgccaccctggccgaactgcgaatgaccggctcggccaccgtcgacctggagctgcatcacgcgctcgccgccgacctgacgctgacggaccgaaatgagcagctcctgaccgcggagcttgccaagacgaactggaagagccaaattctgcttgccctggacctgaacaagacgggcatcacggtgaaggagtacttctatccggcactcaaagccgcggcgacgggccactccgtcgcggagctgtgcttttctgctatccgcaaggtggacgtgcagggccgactcgcggcgccatgtaaagcgatcgaagcccacatgcagcggcagacccagacagacatccacttcctgtctgtagacctggtcgagcccggcacgacgcgcttcaaactgtacctgatggaattggaggtgacgctcgccaagctcgaggagcactggacgctcggaggaacgctcaccgacaaggagacgatgcgtggtctgcagatgatccgcgagctatgggtcgaccttgaaatcgtcgacggcaagcgggctgaaccccagcggccatcgctccccggtgaccctctgagcattgtgccgttctttatgaactacgagataaccccggggcagcctctgcccaagcctaaattctacttccccctcatcggcatccccgagctcaagatcgcgaatgtgctggcagccttcttcgagcgtcatggcatgcatgacctggctagggtctacccggagaatctgcaatcatattaccctggcgaggacttggctattgccacagaccgccaggcatggttgtctatctcgtacaccgaggagaaagggccgtacctgaccatgtactaccactaggaattc |
| *Cs*PT3 | atggtattttcaagcgtgtgctcatttccgtcatccctcggtacaaatttcaaactcgtccctcgttctaattttaaggccagctcgtcgcactatcacgaaattaacaattttatcaacaataaaccgattaagttcagttatttcagctcccgcctgtactgtagtgctaagccgattgttcaccgtgaaaacaaattcaccaaatcgttctcactgagccacctgcagcgcaagagcagcattaaggcgcatggtgagattgaagcggatggcagtaatggtacctctgagtttaatgttatgaaaagcggcaatgcgatctggaggtttgtgcgcccgtatgccgccaagggagtgctgttcaacagcgccgccatgttcgctaaggaactggtgggaaatctaaacctattttcatggccgctaatgtttaaaatcctgtcattcacgctggttattctttgcatttttgtctccacatcaggcattaaccagatttatgatcttgacatcgaccggttgaacaaacctaatctgccggttgcctcaggtgaaatttcagtggaattagcgtggctgctgactattgtgtgcaccatcagcggtttaacgttaacgattattaccaactcaggaccgttctttccatttctgtattccgcatctatcttctttgggtttctgtacagcgctcctccctttcgatggaagaaaaatcccttcaccgcgtgcttctgcaacgtaatgctgtacgtcggaacctcggtcggagtgtactacgcttgtaaagccagccttggcctgccagccaattggtcgccggcattttgcctgctgttttggtttatcagtctcttatcgattcccatctcaatcgcaaaagatttatctgacattgaaggtgatcgcaaatttggcatcattacgttttcgacaaaatttggggcaaaaccgattgcctacatctgccacggcctgatgttgcttaactacgtgtcggttatggcggccgcgatcatttggccccagttcttcaatagctcagtgattcttttatcacacgcattcatggcaatttgggtgttgtaccaggcatggattctggaaaaatctaactatgcaacggaaacttgccaaaaatactacatttttctctggattatttttagtcttgaacacgcattctatctcttcatgtaa |
| CoAnaPT | atgagcccactgagcatgcagacagatagcgttcaggggaccgcagaaaataaatcactggaaactaacggtacgtctaacgatcagcagcttccgtggaaagtgctgggaaaaagcttgggcttaccaaccattgaacaggaacaatactggctgaataccgcgccttacttcaacaatctgctgatccaatgtggctacgatgttcaccaacagtaccagtaccttgctttctaccaccgccacgttctgccggttctgggcccgtttatccgtagctctgctgaagccaactacatctctggcttcagcgcggaaggttacccaatggaactgagcgtgaactatcaggccagtaaagcaacagtacgcctggggtgcgaacccgtgggggagttcgcgggcacgtcacaagatcctatgaaccagttcatgacccgtgaagttcttggccggttgtcccgcctagatcctacgttcgatctgcggctgtttgactatttcgacagtcaattctccctgaccacgtcggaagccaatttggcagcctctaaacttattaaacaacgacgccagagtaaggtcatcgcgttcgacctgaaggatggggcaattattcctaaagcgtatttctttctgaaaggtaaatctttagcaagcggcataccggtgcaggatgttgcttttaatgcgatcgaatccatcgctccgaaacagatcgaatcgccgctgcgtgtcttgcgaaccttcgttaccaagttgttcagcaagccgaccgttacctctgatgtttttatactcgcagtagactgtattgtaccggaaaaatcacgcattaaactgtatgtagcggattcgcagctgtccttggccacattacgcgaattttggacccttgggggtagcgtgactgatagtgcgactatgaaagggctggaaattgcggaggaattgtggcggattctgcagtatgatgatgcggtgtgcagtcatagtaacatggatcaactgccgctggtggtcaattacgaactgtcgtcagggagcgccaccccgaaaccgcagttgtatttgccgttgcacggtcgcaacgacgaagcgatggccaatgctcttaccaaattttgggactatttggggtggaaagggctggctgctcaatataaaaaggacttgtacgcaaacaacccgtgccgcaatctggccgagacaacgacggttcagcggtgggtcgccttcagctacaccgaatcaggcggcgcctaccttacagtgtactttcacgccgttggaggcatgaaaggtaacctgtaa |
| CloQ | atgccggctttacctatagatcaggaatttgactgcgaacgctttcgcgcggatatcagggccacagcggctgcgatcggtgcacctattgcccatcgtctgaccgataccgtcttggaggcgtttcgtgacaattttgcccaaggagccaccctgtggaaaaccacttcacaaccgggtgatcagctgagctatcggttcttcagtcgtctgaagatggacacagtctcacgcgcgattgacgcgggcctgcttgatgcggcccaccccaccttggccgtggtagatgcatggtcatccttgtatggcggtgctccggtgcaaagcggtgattttgatgcaggcagaggtatggcaaaaacgtggctgtatttcggagggctccgcccagcagaggatattctaacagttccagcattacccgcaagcgtacaggcgcgtttgaaagattttctcgctttgggattagcgcatgtacggttcgcggcagtggattggcggcaccattcggcgaacgtatacttccgtgggaagggtcccctggataccgtccaatttgcgcgtatccatgcactaagcggcagcaccccgcctgcggcccatgtggtggaagaagtgttagcttatatgccagaagattactgcgtggcgattacccttgacctgcatagcggagatatcgaacgcgtttgtttttacgcgcttaaagtgccgaagaatgcgctaccacgtatccctactcgcatcgcacggttcttggaggttgcgccgtctcatgatgttgaggagtgtaatgtcattggctggagtttcgggcgctccggcgactatgtaaaagcagagcgctcctacacgggtaatatggcggaaatcctggcgggttggaattgctttttccacggagaagaaggtcgtgatcacgacctccgtgcactgcaccaacacaccgaatctaccatgggcggtgcgcgttgattgtagact |
| *Ec*PT | atggattttccgcagcagttggaagcgtgcgttaaacaagcaaaccaggccctgtctcgttttatcgcgccgcttccgtttcagaatactccggtggttgaaactatgcagtacggcgcgcttctcggagggaaacgtctgcgaccgtttctggtttacgcgaccggtcatatgtttggtgtttctaccaacactctggatgcgcccgcggcggcagtggaatgcattcacgcctactcgctgattcacgacgatctgccggccatggatgacgatgacttgcgtcgaggattacctacttgccacgtaaagttcggtgaagccaatgcaattcttgctggtgatgctctgcaaacgctagcttttagcatactctctgatgccgatatgccggaagtaagtgaccgtgatcgaatctccatgataagcgaactggcatcggcgagcggaatcgcgggtatgtgcggtggccaggccttggacttggatgcagaaggaaaacatgttcccttagacgcacttgagcgaatccaccgtcataaaactggcgccttgattcgcgcagccgtgcgcctgggcgcgctttccgcgggggataagggccgtcgcgccctccctgtgttagataaatatgcggaatccattggtctggcgtttcaggtccaggacgacattttggatgtcgtcggcgataccgcgaccttaggtaaacgccagggcgccgatcagcaactcggtaaatcgacctatcccgcactgctgggcctggaacaagctcgaaagaaagcccgcgatctgattgatgatgcgcgacagagccttaagcagctggcggaacagtccctggatacttctgccctcgaagccctggcggactatattatccaacgaaataaataattgtag |
| NphB | atgtctgaagccgcagatgtagagcgcgtttacgccgcgatggaagaagcggcgggtttgcttggggtggcttgtgcccgggataaaatttacccgcttctcagcacctttcaagacacacttgttgaaggcggctcggtcgttgtcttctcaatggcgagcggccgtcacagcacagaactcgatttctcaatttcggtcccaacaagccacggcgatccgtacgcaactgttgtagaaaagggtctgtttccagcaactgggcacccggtggatgatctgcttgccgacacgcaaaaacacctaccggtcagtatgtttgcaatcgacggcgaagtgaccggtggttttaaaaagacgtacgcgtttttccccaccgacaacatgccgggtgtggcagaactcagcgctattccgagcatgccgccagcggtggcggaaaacgccgaactgtttgcgcgctacggactggataaagtgcagatgacgagcatggattacaagaaacgtcaggtcaatctgtacttttcagaacttagcgcccaaaccctggaagcagaaagcgtactggctcttgtccgcgaactgggtctgcatgtgccgaacgaattaggccttaaattttgtaaacgcagctttagcgtctaccctacgctgaactgggaaaccggcaaaattgatagactgtgcttcgcggtcattagtaacgatcctaccctagtgccctctagtgacgaaggcgatattgaaaaatttcacaactacgcgaccaaggcaccatacgcctacgtgggtgaaaaacggaccctggtgtatggtttaacgttgagccctaaagaagaatactataaactgggtgcgtactatcatatcactgatgtacaacgtggtctgctgaaagcctttgacagcttagaggattga |
| *Sp*PT | atgttggagggaggaatgtctatgagcgaagcagcagatgtggaacgcgtgtacgcggcgatggaagaggccgcgggtttgttgggcgtggcatgcgcgcgcgataaaatctatcctttactttctacatttcaggataccctggtagagggcggctcagtcgtcgtattctcgatggcgtcgggccgtcactcaacggaattggatttttctatctctgtgccgaccagccatggtgatccttatgcgaccgttgtagaaaaaggcctgtttcccgcgacgggtcatccggtggatgacctgttagcggatactcagaaacatctgccggtgtcgatgtttgcaatcgatggcgaagtgacagggggcttcaaaaagacctacgcgttcttcccgacagataacatgccgggtgtggcggagctcagtgctattccaagcatgcccccggcggtagccgaaaatgcagaacttttcgcacgttacgggctggacaaagttcagatgacttcgatggattacaagaaacgtcaggtaaacttatatttttctgagctatctgcacagacgttggaagcggagagcgtgcttgcgctggtgcgcgagctgggtctgcacgtgccaaatgaactgggcttgaagttttgcaaacgtagcttctccgtgtaccccactctgaactgggaaaccggtaaaatcgatcgcctgtgttttgccgtgatcagtaatgatccgaccctcgtcccttcatcggatgaaggcgatattgagaagttccataactacgcaactaaagcgccatatgcctatgtcggcgagaaacgtaccctcgtttatgggctgaccctgagcccgaaagaagagtattataaactgggtgcatactatcatattaccgatgttcagcgtggtctgctgaaggccttcgacagcttagaagactaa |
| UbiA | atggaatggagcttaacccagaataagttactggcgttccatcgtcttatgcgtacagataaaccgattggtgcgttactgttattatggcctaccctctgggccttatgggtcgcgacccccggcgtcccgcagttgtggattctcgccgtttttgtagcgggagtatggctgatgcgtgcagccggctgcgtcgtgaatgattatgccgatcgcaaattcgatggacacgtgaaacgtacggcgaatcgccccctgccttcaggcgcggttaccgagaaagaagcgcgcgccctgtttgtggtcttggtgctgatttcgtttcttttggtgctgacccttaatacgatgaccattctgctgtccattgcggcgcttgccttagcgtgggtctacccgtttatgaaacgctatacccatctgccgcaagtggtcctgggcgcggcgttcggttggtctattccaatggccttcgccgcagtaagcgaaagtgtgccgctgtcgtgctggctgatgtttctggccaacattttatgggcagttgcctacgatacccagtacgctatggtagatcgggatgacgatgttaagattggcatcaaaagcaccgcgattctgttcggccagtatgataaattaattatcggcattctccaaatcggcgtgctggccctcatggcaatcattggggaactgaacggcttgggctggggttattactggagcatcctggtggccggcgcactgttcgtatatcagcaaaagcttattgccaatcgcgaacgtgaagcctgcttcaaagccttcatgaataacaattacgtgggtctggtgttatttctggggttggccatgagttactggcacttcta |

**Table S4. List of primers used for the construction and verification of pathway plasmids.**

| **Primer** | **Sequence^1^** | **Restriction Enzyme** |
| --- | --- | --- |
| NARinsert-Fwd | **AACGTCGTCGCTAAC**GCGACTCCTGCATTAGGAAA | - |
| NARinsert-Rev | CTTTACCAGACTCGAGTCAG | - |
| pRSFDuet-*Fj*TAL-*Cm*CHS-Fwd | TCGAGTCTGGTAAAGAAACCGC | - |
| pRSFDuet-*Fj*TAL-*Cm*CHS-Rev | GTTAGCGACGACGTTTCGGC | - |
| *Hl*PT-Fwd | AAAAAGGATCCG*ATG*GAATTGTCATCCGTG | *Bam*HI |
| *Hl*PT-Rev | AAAAAGAATTC*TCA*TATAAAGAGGTACACTACA | *Eco*RI |
| *Sf* N8DT-1-Fwd | AAAAAGGATCCG*ATG*GGTTCAATGCTGTTAGC | *Bam*HI |
| *Sf* N8DT-1-Rev | AAAAAGAATTC*TTA*GCGGAACAGCGGGA | *Eco*RI |
| AnaPT-Fwd | AAAAAGAGCTCG*ATG*CCTCCCTTGTCTATG | *Sac*I |
| AnaPT-Rev | AAAAAAAGCTT*CTA*GAGATTGCCCTTCAT | *Hind*III |
| CdpC3PT-Fwd | AAAAAGGATCCG*ATG*ACAGTGTCGTCGAC | *Bam*HI |
| CdpC3PT-Rev | AAAAAGAATTC*CTA*GTGGTAGTACATGGTC | *Eco*RI |
| pCDFDuet-Fwd | GAATTCGAGCTCGGC | - |
| pCDFDuet-Rev | CGGATCCTGGCTGTG | - |
| *Cs*PT3-Fwd | **CACAGCCAGGATCCG***ATG*GTATTTTCAAGCGTG | - |
| *Cs*PT3-Rev | **GCCGAGCTCGAATTC***TTA*CATGAAGAGATAGAATG | - |
| coAnaPT-Fwd | **CACAGCCAGGATCCG***ATG*AGCCCACTGAGC | - |
| coAnaPT-Rev | **GCCGAGCTCGAATTC***TTA*CAGGTTACCTTTCATGC | - |
| CloQ-Fwd | **CACAGCCAGGATCCG***ATG*CCGGCTTTACCTATAG | - |
| CloQ-Rev | **GCCGAGCTCGAATTC***AGT*CTACAATCAACGCGC | - |
| *Ec*PT-Fwd | **CACAGCCAGGATCCG***ATG*GATTTTCCGCAGCAG | - |
| *Ec*PT-Rev | **GCCGAGCTCGAATTC***CTA*CAATTATTTATTTCGTTGGA | - |
| NphB-Fwd | **CACAGCCAGGATCCG***ATG*TCTGAAGCCGCAG | - |
| NphB-Rev | **GCCGAGCTCGAATTC***TCA*ATCCTCTAAGCTGTCAA | - |
| *Sp*PT-Fwd | **CACAGCCAGGATCCG***ATG*TTGGAGGGAGGAAT | - |
| *Sp*PT-Rev | **GCCGAGCTCGAATTC***TTA*GTCTTCTAAGCTGTCG | - |
| UbiA-Fwd | **CACAGCCAGGATCCG***ATG*GAATGGAGCTTAAC | - |
| UbiA-Rev | **GCCGAGCTCGAATTC***TAG*AAGTGCCAGTAACTC | - |
| MCS1-Fwd | GGATCTCGACGCTCTCC | - |
| *Cm*CHS-sequencing-Fwd | CCGAAACGTCGTCGCTAA | - |
| T7terminator-Rev | CTAGTTATTGCTCAGCGGT | - |

^1^ Start and stop codons in *italic*; restriction enzymes are underlined; homology arms in **bold**. To maintain the sequence in frame, one base was occasionally added between the restriction site and the gene start codon.

**Table S5. Sequence of the genes used in the genome integration strategies.**

| **Gene** | **Sequence** |
| --- | --- |
| *Ec*DXS | atgagttttgatattgccaaatacccgaccctggcactggtcgactccacccaggagttacgactgttgccgaaagagagtttaccgaaactctgcgacgaactgcgccgctatttactcgacagcgtgagccgttccagcgggcacttcgcctccgggctgggcacggtcgaactgaccgtggcgctgcactatgtctacaacaccccgtttgaccaattgatttgggatgtggggcatcaggcttatccgcataaaattttgaccggacgccgcgacaaaatcggcaccatccgtcagaaaggcggtctgcacccgttcccgtggcgcggcgaaagcgaatatgacgtattaagcgtcgggcattcatcaacctccatcagtgccggaattggtattgcggttgctgccgaaaaagaaggcaaaaatcgccgcaccgtctgtgtcattggcgatggcgcgattaccgcaggcatggcgtttgaagcgatgaatcacgcgggcgatatccgtcctgatatgctggtgattctcaacgacaatgaaatgtcgatttccgaaaatgtcggcgcgctcaacaaccatctggcacagctgctttccggtaagctttactcttcactgcgcgaaggcgggaaaaaagttttctctggcgtgccgccaattaaagagctgctcaaacgcaccgaagaacatattaaaggcatggtagtgcctggcacgttgtttgaagagctgggctttaactacatcggcccggtggacggtcacgatgtgctggggcttatcaccacgctaaagaacatgcgcgacctgaaaggcccgcagttcctgcatatcatgaccaaaaaaggtcgtggttatgaaccggcagaaaaagacccgatcactttccacgccgtgcctaaatttgatccctccagcggttgtttgccgaaaagtagcggcggtttgccgagctattcaaaaatctttggcgactggttgtgcgaaacggcagcgaaagacaacaagctgatggcgattactccggcgatgcgtgaaggttccggcatggtcgagttttcacgtaaattcccggatcgctacttcgacgtggcaattgccgagcaacacgcggtgacctttgctgcgggtctggcgattggtgggtacaaacccattgtcgcgatttactccactttcctgcaacgcgcctatgatcaggtgctgcatgacgtggcgattcaaaagcttccggtcctgttcgccatcgaccgcgcgggcattgttggtgctgacggtcaaacccatcagggtgcttttgatctctcttacctgcgctgcataccggaaatggtcattatgaccccgagcgatgaaaacgaatgtcgccagatgctctataccggctatcactataacgatggcccgtcagcggtgcgctacccgcgtggcaacgcggtcggcgtggaactgacgccgctggaaaaactaccaattggcaaaggcattgtgaagcgtcgtggcgagaaactggcgatccttaactttggtacgctgatgccagaagcggcgaaagtcgccgaatcgctgaacgccacgctggtcgatatgcgttttgtgaaaccgcttgatgaagcgttaattctggaaatggccgccagccatgaagcgctggtcaccgtagaagaaaacgccattatgggcggcgcaggcagcggcgtgaacgaagtgctgatggcccatcgtaaaccagtacccgtgctgaacattggcctgccggacttctttattccgcaaggaactcaggaagaaatgcgcgccgaactcggcctcgatgccgctggtatggaagccaaaatcaaggcctggctggcataa |
| *Ec*IDI | atgcaaacggaacacgtcattttattgaatgcacagggagttcccacgggtacgctggaaaagtatgccgcacacacggcagacacccgcttacatctcgcgttctccagttggctgtttaatgccaaaggacaattattagttacccgccgcgcactgagcaaaaaagcatggcctggcgtgtggactaactcggtttgtgggcacccacaactgggagaaagcaacgaagacgcagtgatccgccgttgccgttatgagcttggcgtggaaattacgcctcctgaatctatctatcctgactttcgctaccgcgccaccgatccgagtggcattgtggaaaatgaagtgtgtccggtatttgccgcacgcaccactagtgcgttacagatcaatgatgatgaagtgatggattatcaatggtgtgatttagcagatgtattacacggtattgatgccacgccgtgggcgttcagtccgtggatggtgatgcaggcgacaaatcgcgaagccagaaaacgattatctgcatttacccagcttaaataa |
| *Bs*DXS | atgtcaattgacgagcttgaaaagctgagcgatgaaatccgccaatttctgatcacaagtttaagcgcgtccggtggtcatattggtcccaatttgggcgttgtcgagttgaccgtagcgcttcataaagaatttaactctccaaaagataaatttctttgggacgtaggccatcagagttacgttcacaaattgttgacaggccgtggtaaagagttcgctactttacgtcaatacaagggattatgtggtttcccgaagcgtagtgagtcggaacacgatgtttgggagacagggcacagtagtacctccttgtcgggagctatgggtatggcagcagcccgtgatattaaaggaactgacgaatacatcattcccattatcggagacggggctcttactggtggcatggctttggaagcacttaatcatatcggggacgaaaaaaaggatatgattgttattttaaacgataatgagatgtccattgctccaaacgtcggggcgatccattcaatgttaggccgtttacgcaccgccggcaagtatcaatgggttaaggacgaactggaatatctgttcaaaaagattcccgcagtcggcgggaaattggctgctactgctgaacgtgtgaaggattctttgaagtacatgttagtgtcaggcatgtttttcgaggaattgggttttacctatttgggtccggttgatggacacagttaccacgaacttatcgaaaacttacagtatgctaagaagacgaaaggccctgttcttcttcacgtgatcacaaaaaaggggaagggatataaacccgctgagacggacacaattggcacgtggcatggcacaggaccatacaaaattaacacaggggacttcgtaaagccgaaagcagcggccccttcttggtccgggttggtcagcggtaccgtacagcgtatggcacgtgaggatgggcgcatcgtcgccattacgccggctatgccagtaggttccaagcttgaaggctttgcgaaggagttccctgaccgtatgttcgatgtaggtatcgccgaacaacatgctgctacgatggccgctgcaatggccatgcaaggcatgaagccatttttggcaatctatagcacgttcttgcaacgtgcatatgaccaagttgtgcatgacatttgtcgtcagaatgctaacgtgttcattggcatcgaccgtgccgggctggtaggagcagatggtgaaacacaccaaggcgtctttgacattgcattcatgcgtcatatccctaacatggtgcttatgatgcctaaggatgagaatgagggacaacacatggtccacacggctttatcttacgatgaaggtccgatcgctatgcgctttcctcgtgggaatggattaggggtaaaaatggacgaacaattaaagacgatccctatcgggacatgggaggttttacgtccaggcaacgacgcggtaatcttgacgttcgggaccacaattgagatggcaattgaggccgccgaggagttgcaaaaagaaggcttatctgtacgcgttgtgaatgctcgtttcatcaagcctattgacgaaaagatgatgaagagcattctgaaagaagggctgccgattttaaccatcgaggaggcggtccttgaaggggggttcgggagctccatcctggagtttgcacacgatcagggagaatatcatactccaattgatcgtatgggtattccagaccgtttcattgaacatggctccgtaacggcgctgctggaggaaattggcttaacaaaacaacaggttgcaaatcgcattcgcttactgatgcctcccaaaacacataaagggattggctcctga |
| *Sc*IDI | atgacggcggataacaatagtatgccacatggtgctgtgtcctcttacgctaaactggtgcagaatcagacgccagaggatatcttagaagaatttccggagatcattcctttacagcaacgtccaaacactcgcagttccgagacgagcaatgacgaatccggggagacctgttttagtggtcacgatgaagagcaaatcaaattaatgaacgaaaactgcattgttctggactgggatgacaatgccattggcgccgggacgaagaaggtgtgtcacttaatggagaacattgagaaagggttgttacaccgtgccttctcggtttttatctttaatgaacagggggaactgctgttgcagcagcgtgcaactgaaaagatcacattcccggatctgtggacgaacacgtgttgctctcacccactttgcatcgacgatgaattaggtttgaaagggaagttagatgataagattaaaggggctattacagcggcagtgcgtaaactggaccacgagttagggattccagaagacgagacgaaaacgcgcgggaaatttcatttccttaatcgtatccactacatggccccgtcgaacgaaccgtggggtgagcacgaaatcgattacatccttttctataaaatcaacgctaaggaaaacctgacggtgaaccccaacgtaaacgaggtacgcgactttaagtgggtaagcccaaacgacttgaaaactatgttcgctgatccctcatataaatttaccccgtggttcaaaattatctgtgaaaactatctttttaactggtgggagcaattagatgacttatccgaggtagaaaacgatcgtcaaatccatcgcatgctttaa |
| *Bl*IDI | atggtgacgcgtgcaaagcgcaagaaagaacatattgaccacgctttaagcacgggtcagaaacgccaaactggtttggatgatatcacgttcgtacacgtatcgcttccagagactgagttgtcacaggtagacacaagtaccaagattggtgagctttttttgtcctcccccattttcattaacgccatgacgggtggagggggaaaggcaaccttcgagatcaatcgtgctttggcccgcgcggccgcgcaaactgggatccctgtggctgtcggttctcagatgtcggcattgaaagatccagatgagcgtcctagctatgagatcgtgcgcaaagagaatatgaagggccttgtattcgcgaatttagggagcgaggcgactgttgagcaggccaaacgcgccgtcgatatgatcgaagctgacatgctgcaaattcaccttaacgtcattcaagagatcgtgatgccggaaggagaccgcaattttaccgggcgtctgcgtcgcatcgaagatatttgccgttcggtgtcagtgccagtggcggtcaaggaggtcggatttgggatgagtcgcgataccgccgcgcgtttatttaatgttggagtccaggccatcgacgtcggcgggttcggagggacaaacttttccaagatcgaaaaccttcgtcgtgataaggccgtagagttcttcgaccagtggggtatctcaacggcggcaagcctggctgaggtatctagtatttctggtgatcgccccattatcgccagtgggggcatccaggacgcattggatcttgcaaagagtatcgcattgggtgctagcgccgctgggatggccggttattttttgaaagtccttacagccagcggggaggaagcattagccgccgaaatcgagtcgcttatcgaggacttcaagcgcatcatgacggtccttgggtgccgcacgattgaacaacttaagaaagccccccttgtcatcaagggggacacttatcattggttgaaggcgcgtggagtagacccgtttgtctattccatgcgttaa |

**Table S6. List of primers used for the construction and verification of CRISPR-Cas12a plasmids for genome integration.**

| **Primer** | **Sequence^1^** |
| --- | --- |
| pTF-integration*lacZ*-Fwd | CTCGAATGGTCTATATCCTACG |
| pTF-integration*lacZ*-Rev | ATGTATATCTCCTTCTTAAAAGATC |
| *Bs*DXS-integration1-Fwd | **GAAGGAGATATACAT***ATG*TCAATTGACGAGCTTGAA |
| *Bs*DXS-integration1-Rev | **TATAGACCATTCGAG***TCA*GGAGCCAATCCCTTTATG |
| *Bs*DXS-integration2-Rev | *TCA*GGAGCCAATCCCTTTATG |
| *Sc*IDI-integration1-Fwd | **GAAGGAGATATACAT***ATG*ACGGCGGATAACAATAGT |
| *Sc*IDI-integration2-Fwd | **GGGATTGGCTCCTGA***ATG*ACGGCGGATAACAATAGT |
| *Sc*IDI-integration1-Rev | **TATAGACCATTCGAG***TTA*AAGCATGCGATGGATTTG |
| *Bl*IDI-integration1-Fwd | **GAAGGAGATATACAT***ATG*GTGACGCGTGCAAAGC |
| *Bl*IDI-integration2-Fwd | **GGGATTGGCTCCTGA***ATG*GTGACGCGTGCAAAGC |
| *Bl*IDI-integration1-Rev | **TATAGACCATTCGAG***TTA*ACGCATGGAATAGACAAAC |
| *Ec*DXS-integration1-Fwd | **GAAGGAGATATACAT***ATG*AGTTTTGATATTGCCAAATACC |
| *Ec*DXS-integration1-Rev | **TATAGACCATTCGAG***TTA*TGCCAGCCAGGCC |
| *Ec*DXS-integration2-Rev | *TTA*TGCCAGCCAGGCC |
| *Ec*IDI-integration1-Fwd | **GAAGGAGATATACAT***ATG*CAAACGGAACACGT |
| *Ec*DI-integration2-Fwd | **GCCTGGCTGGCATAA***ATG*CAAACGGAACACGT |
| *Ec*IDI-integration1-Rev | **TATAGACCATTCGAG***TTA*TTTAAGCTGGGTAAATGC |
| pTF-sequencing-Fwd | gcgtgactacctacgggtaa |
| pTF-sequencing-Rev | gcgttacgcgttcgctca |

^1^ Start and stop codons in *italic*; homology arms in **bold**.

**Table S7. Sequence of geranyl diphosphate/farnesyl diphosphate synthase (*ispA*) gene.**

| **Gene** | **Sequence^1^** |
| --- | --- |
| *ispA* | atggac**tttc**cgcagcaactcgaagcctgcgttaagcaggccaaccaggcgctgagccgttttatcgccccactgccctttcagaacactcccgtggtcgaaaccatgcagtatggcgcattattaggtggtaagcgcctgcgacctttcctgg**ttta**tgccaccggtcatatgttcggcgttagcacaaacacgctggacgcacccgctgccgccgttgagtgtatccacgcttactcattaattcatgatgatttaccggcaatggatgatgacgatctgcgtcgcggtttgccaacctgccatgtgaagtttggcgaagcaaacgcgattctcgctggcgacgctttacaaacgctggcgttctcgattttaagcgatgccgatatgccggaagtgtcggaccgcgacagaatttcgatgatttctgaactggcgagcgccagtggtattgccggaatgtgcggtggtcaggcattagatttagacgcggaaggcaaacacgtacctctggacgcgcttgagcgtattcatcgtcataaaaccggcgcattgattcgcgccgccgttcgccttggtgcattaagcgccggagataaaggacgtcgtgctctgccggtactcgacaagtatgcagagagcatcggccttgccttccaggttcaggatgacatcctggatgtggtgggagatactgcaacgttgggaaaacgccagggtgccgaccagcaacttggtaaaagtacctaccctgcacttctgggtcttgagcaagcccggaagaaagcccgggatctgatcgacgatgcccgtcagtcgctgaaacaactggctgaacagtcactcgatacctcggcactggaagcgctagcggactacatcatccagcgtaataaataa |

^1^ PAM sequences are represented in **bold** and 23 bp array sequences are underlined.

**Table S8. List of primers used for the construction and verification of CRISPRi plasmids.**

| **Primer** | **Sequence^1^** |
| --- | --- |
| pddcpf1-crRNAins-*ispA*1-Fwd | TCTACTCTTGTAGAT**CGCAGCAACTCGAAGCCTGCGTT**TTTTTTTGAAGCTTGGGCCCG |
| pddcpf1-crRNAins-*ispA*2-Fwd | TCTACTCTTGTAGAT**TGCCACCGGTCATATGTTCGGCG**TTTTTTTGAAGCTTGGGCCCG |
| pddcpf1-crRNAins-Rev | ATCTACAAGAGTAGAAATTGAAATTGTTATCCGC |
| pddcpf1-sequencing-Fwd | GCTCGCGTATATTCAGGAACT |
| pddcpf1-sequencing-Rev | AGGCCCACCCGAAGGTGAGCC |

^1^ 23 bp array sequences in **bold**.

**Table S9. List of primers used for reverse transcription-quantitative polymerase chain reaction (RT-qPCR).**

| **Primer** | **Sequence** |
| --- | --- |
| *ispA*-qPCR-Fwd | GTGGTATTGCCGGAATGTGC |
| *ispA*-qPCR-Rev | AATGCGCCGGTTTTATGACG |
| *hcaT*-Fwd | CCGTTTACAGGCGGTTTACT |
| *hcaT*-Rev | CCGTGGCCCAGATATTGATAC |
| *idnT*-Fwd | GTGCGCCTCTTCTTTGAATTT |
| idnT-Rev | TCGATGGTGCGTCCATTAC |

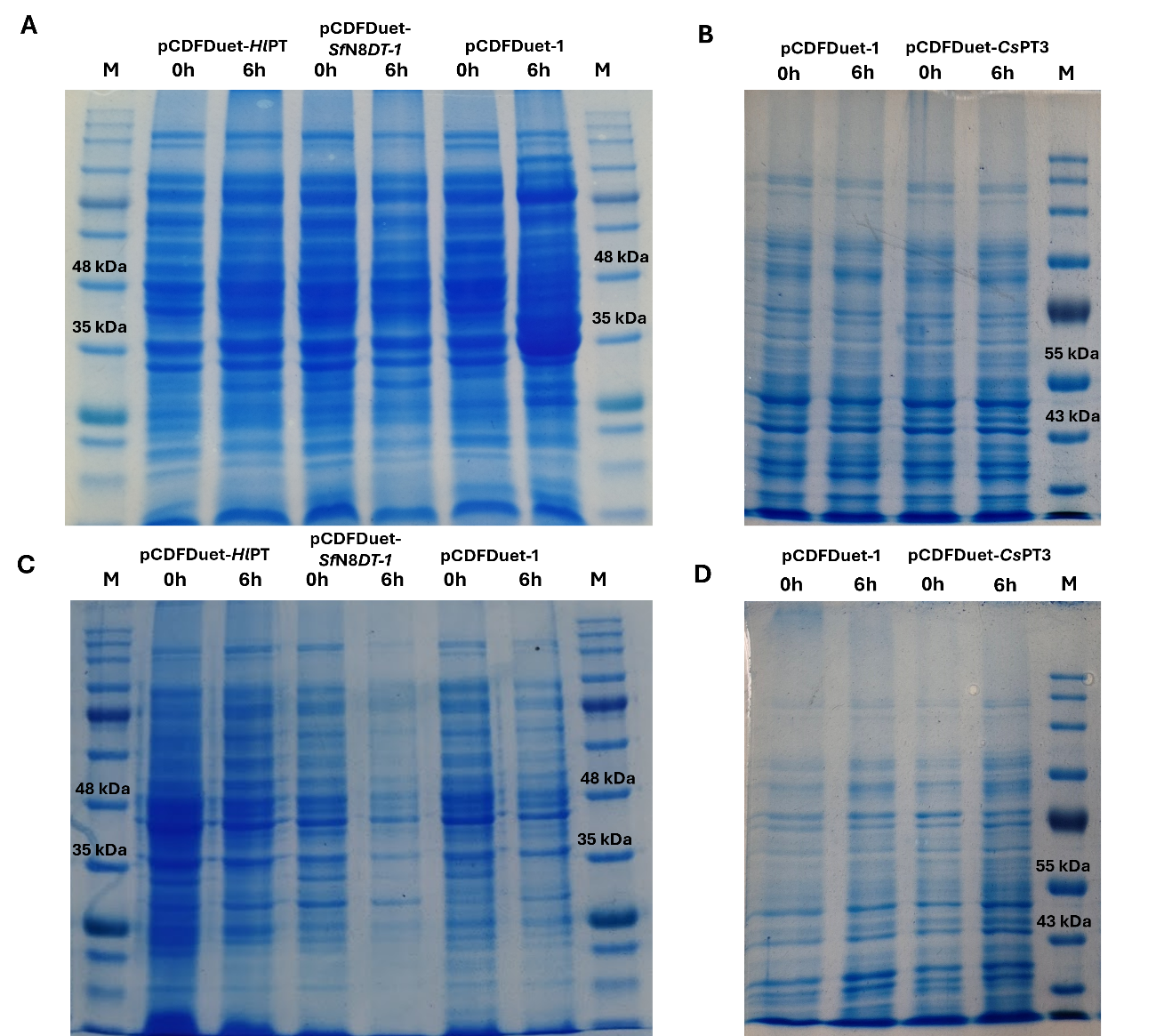

**Figure S1. Protein sodium dodecyl sulfate (SDS) polyacrylamide gel electrophoresis (SDS-PAGE) gels of soluble (A and B) and insoluble (C and D) protein fractions showing plant aromatic prenyltransferases (PTs) expression in *E. coli* M-PAR-121 strain at time zero (0 h) of induction and after 6 h of induction.** PT from *Humulus lupulus* (*Hl*PT), N8DT-1 from Sophora flavescens (SfN8DT-1), and PT3 from *Cannabis sativa* (*Cs*PT3) protein bands are expected to be around 47.29 kDa, 46.84 kDa, and 46.76 kDa, respectively. *E. coli* M-PAR-121 carrying pCDFDuet-1 was used as a control strain. NZYColour Protein Marker II (NZYTech) was used as protein marker in gels A and C. Blue Prestained Protein Standard, Broad Range (11-250 kDa) was used as protein marker in gels B and D.

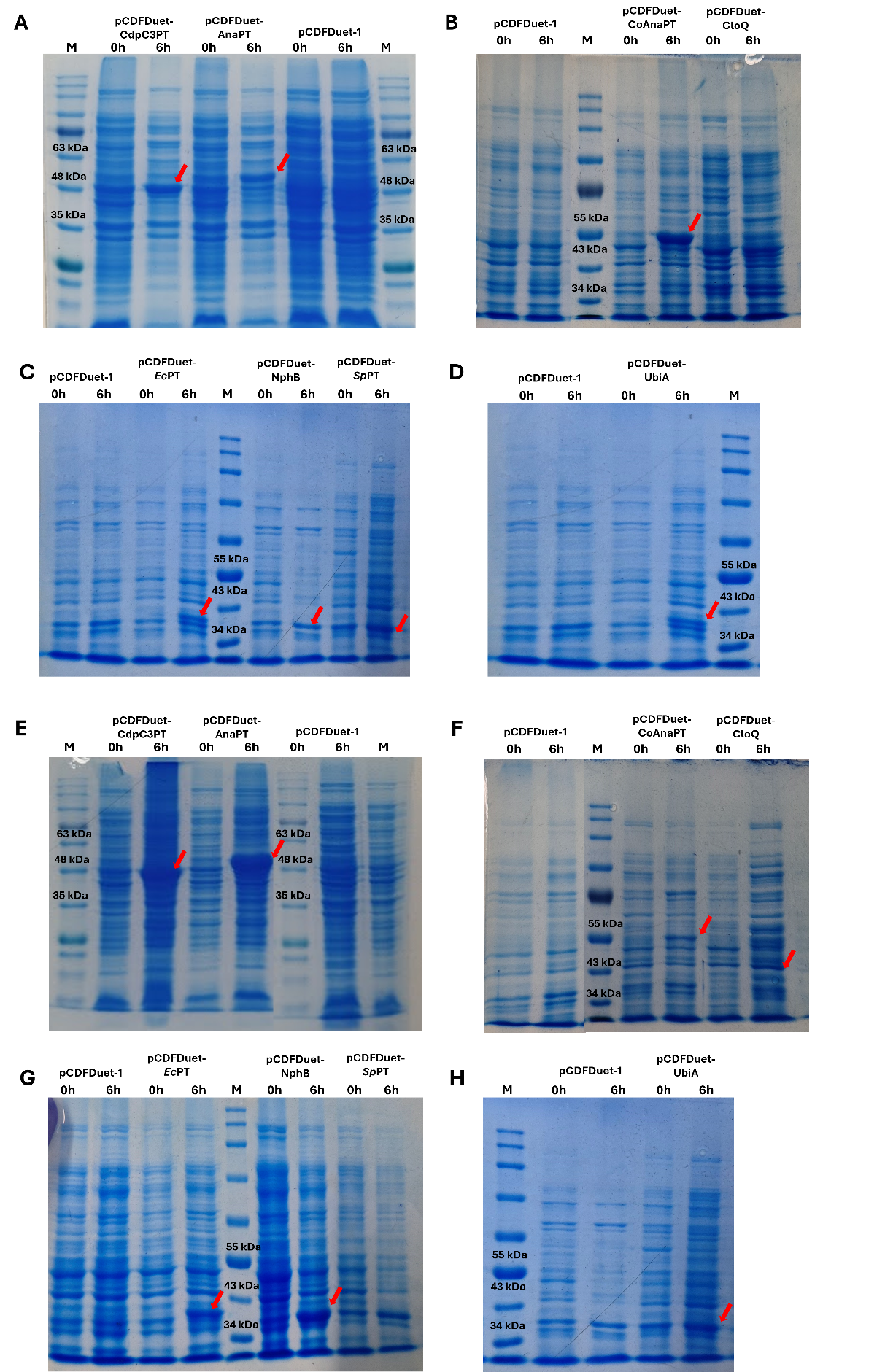

**Figure S2. Protein sodium dodecyl sulfate (SDS) polyacrylamide gel electrophoresis (SDS-PAGE) gels of soluble (A, B, C, and D) and insoluble (E, F, G, and H) protein fractions showing aromatic prenyltransferases (PTs) from microbial sources expression in *E. coli* M-PAR-121 strain at time zero (0 h) of induction and after 6 h of induction.** CdpC3PT and AnaPT from *Neosartorya fischeri* (without codon-optimization) protein bands are expected to be around 48.62 kDa and 50.62 kDa. Codon-optimized AnaPT (coAnaPT), CloQ from *Streptomyces roseochromogenes,* PT from *E. coli (Ec*PT*),* NphB from *Streptomyces* sp., PT from *Streptomyces* sp. (*Sp*PT), and UbiA from *E. coli* were expected at 46.76 kDa, 50.42 kDa, 37.20 kDa, 34.80 kDa, 35.40 kDa, 36.06 kDa, and 34.13 kDa, respectively. *E. coli* M-PAR-121 carrying pCDFDuet-1 was used as a control strain. NZYColour Protein Marker II (NZYTech) was used as protein marker in gels A and E. Blue Prestained Protein Standard, Broad Range (11-250 kDa) was used as protein marker in gels B, C, D, F, G, and H.

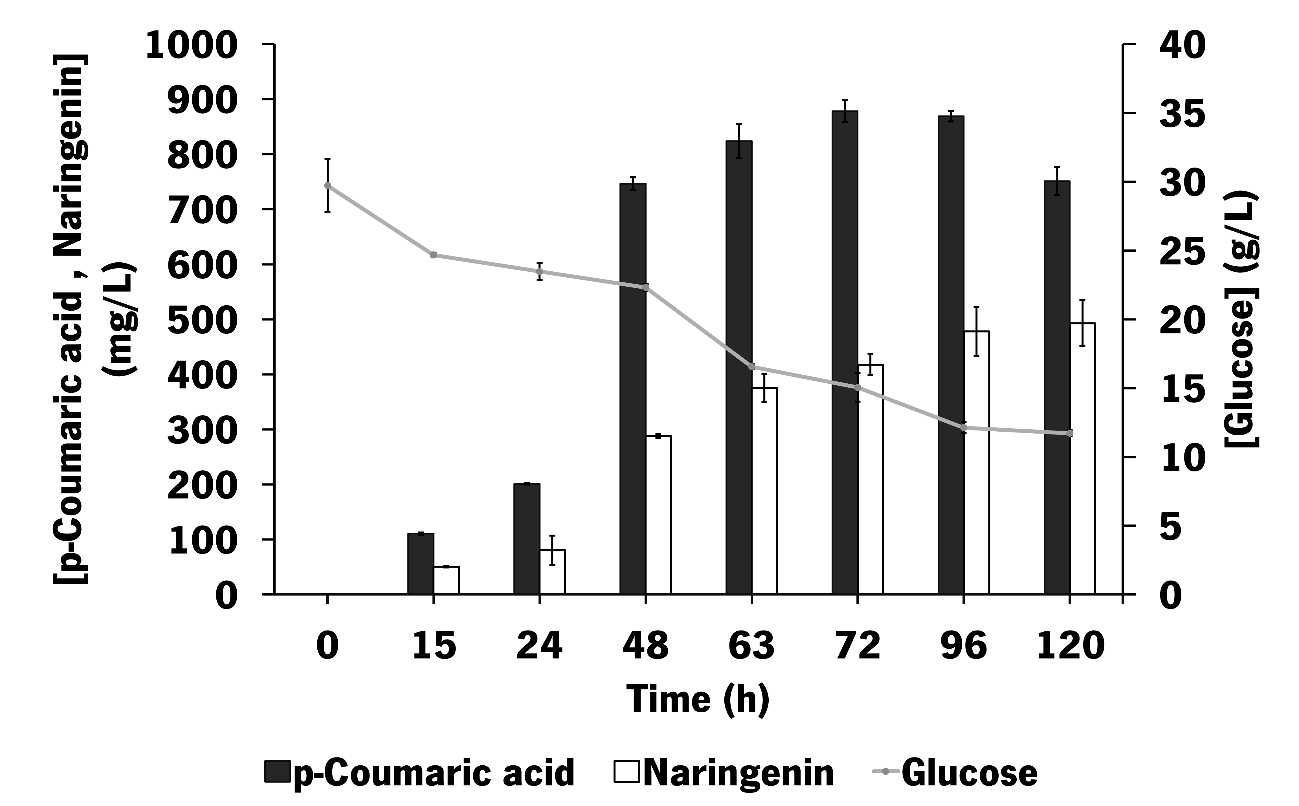

**Figure S3. Profile of glucose consumption and metabolites production by *Escherichia coli* M-PAR-121 expressing pRSFDuet_*Fj*TAL_*Cm*CHS_*At*4CL_*Ms*CHI in shake flask experiments using the combination of LB+M9.** Results correspond to the average of three independent experiments ± standard deviation.

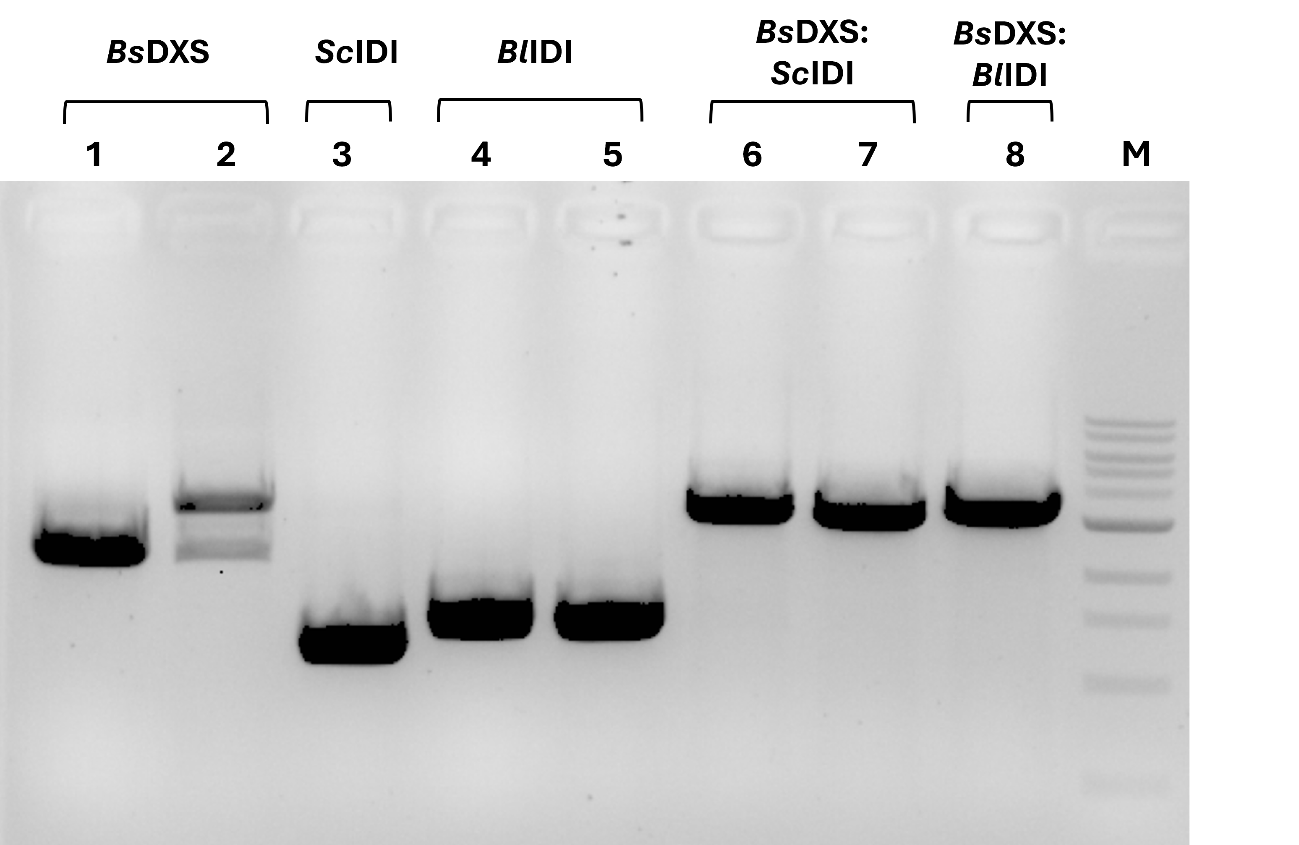

**Figure S4. Agarose gel 1% (w/v) to confirm the correct integration of heterologous 1-deoxy-D-xylulose-5-phosphate synthase (DXS) and isopentenyl diphosphate isomerase (IDI) genes into the *lacZ* locus of *E. coli* M-PAR-121.** The genomic DNA of two colonies was tested for the integration of *Bs*DXS (Lanes 1 and 2), *Sc*IDI (Lanes 3), *Bl*IDI (Lanes 4 and 5), *Bs*DXS:*Sc*IDI (Lanes 6 and 7), and *Bs*DXS:*Bl*IDI (Lane 8). The expected band sizes for the integration of *Bs*DXS, *Sc*IDI, *Bl*IDI, *Bs*DXS:*Sc*IDI, and *Bs*DXS:*Bl*IDI are 2178 bp, 1185 bp, 1371 bp, 3045 bp, and 3231 bp, respectively. The expected size if the integration did not occur (correspondent to a lacZ region) is 575 bp. Transformants 1, 3, 4, 7 and 8 were selected. M corresponds to 1 kb DNA ladder (NEB).

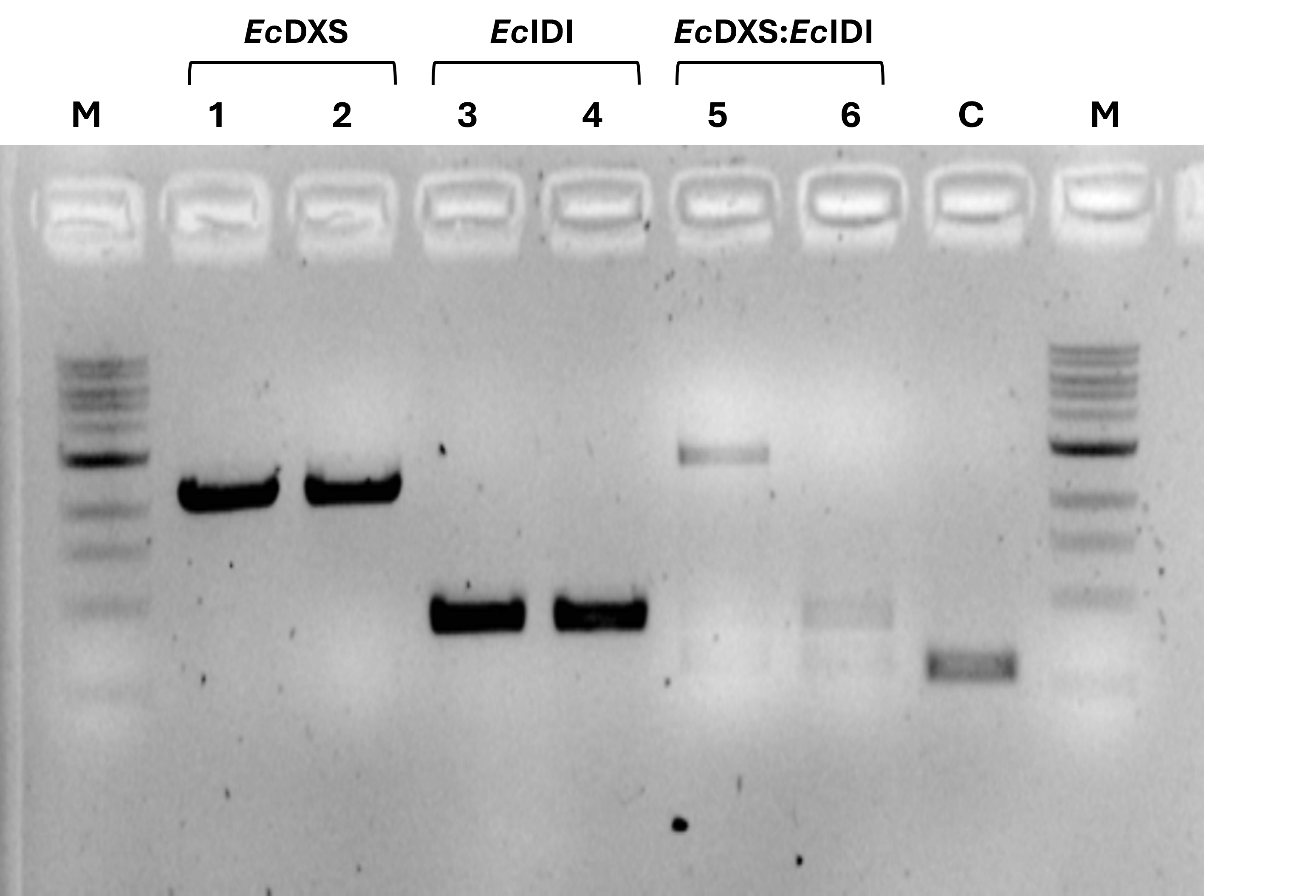

**Figure S5. Agarose gel 1% (w/v) to confirm the correct integration of native 1-deoxy-D-xylulose-5-phosphate synthase (DXS) and isopentenyl diphosphate isomerase (IDI) genes into the *lacZ* locus of *E. coli* M-PAR-121.** The genomic DNA of two colonies was tested for each integration. The expected band sizes for the integration of *Ec*DXS, *Ec*IDI, and *Ec*DXS:*Ec*IDI are 2181 bp, 867 bp, 2730 bp. A control PCR with the genomic DNA of the wild-type strain was performed (Lane C). The expected size for this fragment is 575 bp. Transformants 1, 3, and 5 were selected. M corresponds to 1 kb DNA ladder (NEB).

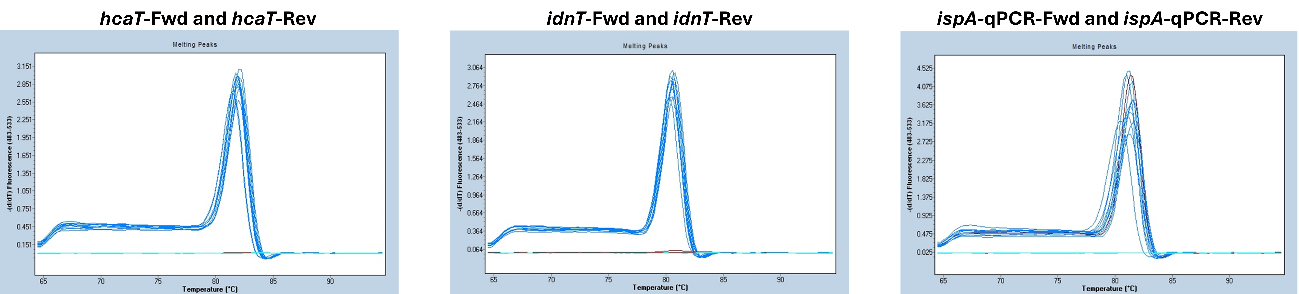

**Figure S6. Melting curves of the primers targeting the housekeeping genes (putative 3-phenylpropionate transporter coding gene (*hcaT*) and gluconate transporter coding gene (*idnT*)) and the geranyl diphosphate/farnesyl diphosphate synthase gene (*ispA*) used in reverse transcription-quantitative polymerase chain reaction (RT-qPCR) for expression levels quantification.**

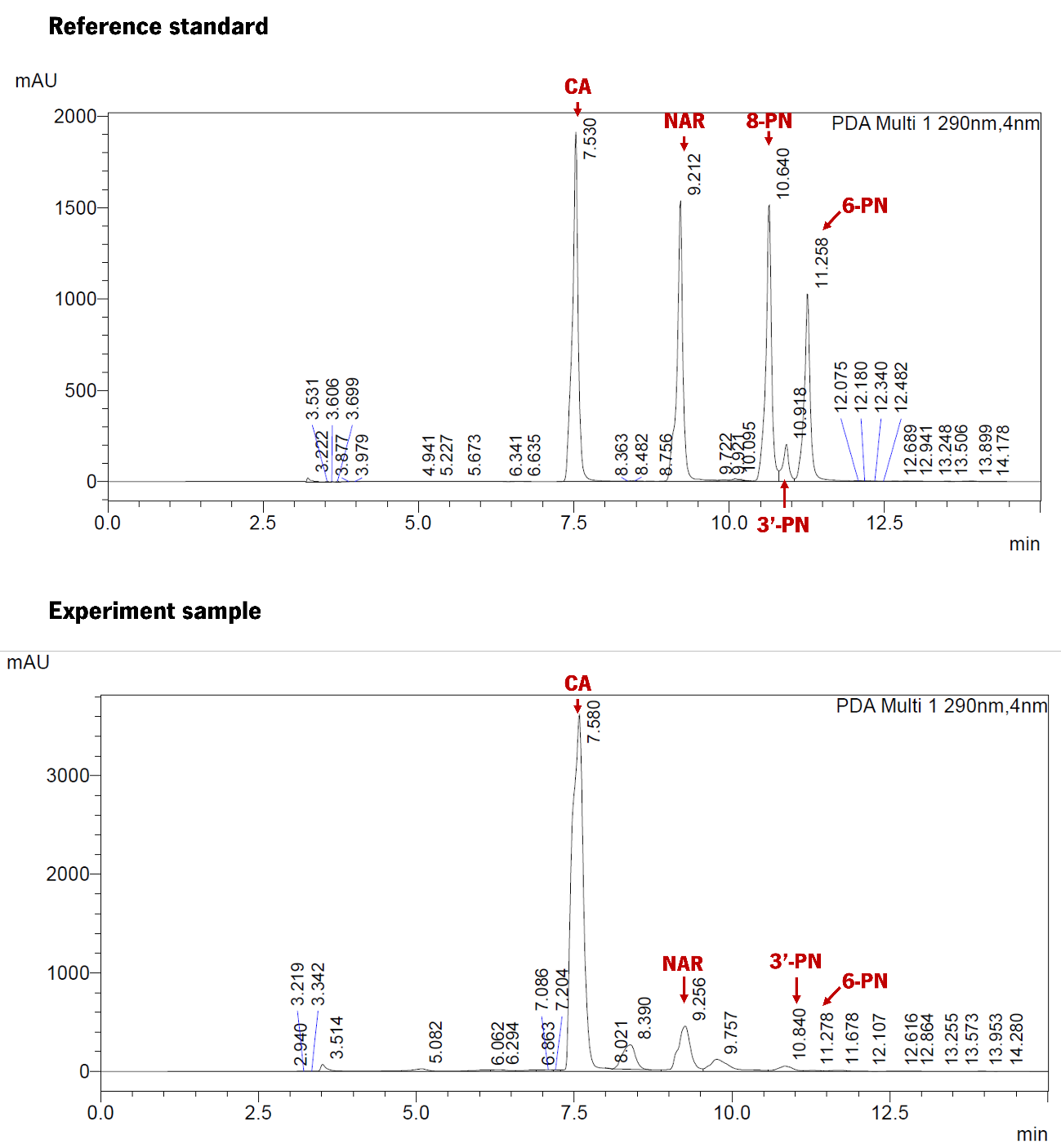

**Figure S7. Representative chromatograms of the analytical standard composed by *p-*coumaric acid (CA), naringenin (NAR), 8-prenylnaringenin (8-PN), 3’-prenylnaringenin (3’-PN) and 6-prenylnaringenin (6-PN) and one sample from the bioreactor experiment with higher production levels.**
